# Evaluation of homologous recombination repair status in metastatic prostate cancer by next-generation sequencing and functional tissue-based immunofluorescence assays

**DOI:** 10.1101/2024.01.28.577367

**Authors:** Sara Arce-Gallego, Pablo Cresta Morgado, Luisa Delgado-Serrano, Sara Simonetti, Macarena Gonzalez, David Marmolejo, Rafael Morales-Barrera, Gisela Mir, Maria Eugenia Semidey, Paula Romero Lozano, Sarai Cordoba-Terreros, Richard Mast, Matias de Albert, Jacques Planas, Mercè Cuadras, Xavier Maldonado, Cristina Suarez, Irene Casanova-Salas, Lara Nonell, Rodrigo Dienstmann, Paolo Nuciforo, Ana Vivancos, Alba Llop-Guevara, Joan Carles, Violeta Serra, Joaquin Mateo

**Affiliations:** Prostate Cancer Translational Research Group, Institute of Oncology (VHIO), Vall d’Hebron University Hospital, Barcelona, Spain; Oncology Data Science, Vall d’Hebron Institute of Oncology (VHIO), Barcelona, Spain; Molecular Pathology Group, Vall d’Hebron Institute of Oncology (VHIO), Barcelona, Spain; Medical Oncology Department, Vall d’Hebron University Hospital and Vall d’Hebron Institute of Oncology (VHIO), Barcelona, Spain; Pathology Department, Vall d’Hebron University Hospital, Barcelona, Spain; Cancer Genomics Group, Vall d’Hebron Institute of Oncology (VHIO), Barcelona, Spain; Department of Radiology, Vall d’Hebron University Hospital, Barcelona, Spain; Urology Department, Vall d’Hebron University Hospital, Barcelona, Spain; Radiotherapy Department, Vall d’Hebron University Hospital, Barcelona, Spain; Bioinformatics Unit, Vall d’Hebron Institute of Oncology (VHIO), Barcelona, Spain; Experimental Therapeutics Group, Vall d’Hebron Institute of Oncology (VHIO), Barcelona, Spain

**Keywords:** prostate cancer, biomarkers, genomics, HRR, BRCA

## Abstract

**Purpose:** Metastatic prostate cancer (mPC) is enriched for homologous recombination repair (HRR) gene alterations; these biomarkers have prognostic and predictive value. Next-generation sequencing (NGS) allows for patient stratification based on these biomarkers, but widespread clinical implementation is still limited. Moreover, not all mutations in HRR genes result in functional HRR loss in the tumor. We investigated the correlation between genomic and functional loss of HRR, using NGS and an optimized RAD51 immunofluorescence (RAD51-IF) assay in mPC clinical biopsies.

**Experimental design:** Observational study including patients with stage IV prostate cancer. Biopsies from either primary tumor or metastatic biopsies underwent NGS (targeted sequencing and/or whole-exome sequencing), and RAD51-IF. Clinical data was extracted from electronic patient records.

**Results:** 219 biopsies from 187 patients were acquired, including primary (151/219) and metastatic (68/219) tumor biopsies collected either in the metastatic hormone-sensitive (169/219) or castration-resistant (50/219) setting. NGS (181 biopsies from 157 patients) showed frequent genomic alterations in *TP53* (40%), *AR* (15%), *PTEN* (14%), MYC (10%), *BRCA2* (9%), *ATM* (8%) and *BRCA1* (2%). Tissue for RAD51 IF was available for 206 samples; of those, 140/206 (68%) were evaluable for RAD51- IF. Based on a previously defined threshold of 10% RAD51-positive cells, 21% samples had RAD51-low results compatible with HRR deficiency (HRD). Sample matched RAD51-IF and genomics data were obtained for 128 biopsies (117 patients): RAD51-IF had a high sensitivity (68%) and specificity (85%) to identify cases with *BRCA1/2* alterations. Additionally, the RAD51-IF assay was able to identify restoration of HRR function in selected cases with *BRCA2* reversion mutations or BRCA1 expression.

**Conclusions:** RAD51-IF is feasible in routine clinical samples from mPC patients and associates strongly with clinically relevant HRR gene alterations.

## INTRODUCTION

Metastatic prostate cancer (mPC) is a lethal disease. Advanced in molecular stratification have led to improved patient outcome, as illustrated by the association between homologous recombination repair (HRR) gene biomarkers (Abeshouse et al., 2015; Abida et al., 2017; Grasso et al., 2012; Robinson et al., 2015), and responses to poly(ADP-ribose) polymerase (PARP) inhibitor treatment (Agarwal et al., 2023; Chi et al., 2023; de Bono et al., 2020; Mateo et al., 2015). Pivotal trials of PARP inhibitors in prostate cancer have shown, however, high inter-patient variability in clinical response among patients with different HRR gene alterations. On one hand, response rates among patients with biallelic *BRCA2* inactivation, arguably the highest predictive biomarkers of PARPi benefit, ranges 45-65% across trials, meaning that almost half of these patients do not achieve PSA or radiological responses. On the other hand, response rates among patients with alterations in non-BRCA HRR genes, or other genes relevant for double-strand break (DSB) repair, are limited. Therefore, biomarkers to complement NGS for accurate patient stratification and treatment selection are needed.

To improve diagnostic and predictive accuracy, the detection of repetitive patterns of DNA fragments or scars resulting from error-prone repair of DSBs in tumors with deficient HRR has been proposed as complimentary to mutation detection. However, the applicability of these biomarkers in prostate cancer management remains controversial, in contrast to their extensive development in breast or ovarian cancer (Davies et al., 2017; Nguyen et al., 2020; Sztupinszki et al., 2020). These include accumulation of loss-of-heterozygosity events (Abkevich et al., 2012), large-scale state transitions (Popova et al., 2012) (LST score), and telomeric allelic imbalance (Birkbak et al., 2012) (TAI score); these three measures can be combined into a single score (Telli et al., 2016) (Davies et al., 2017; Frampton et al., 2013; Nguyen et al., 2020).

One added caveat of next-generation sequencing for treatment selection in mPC arises from the many technical challenges to deliver genomics assays on small prostate biopsies. Different studies reported a 20-40% failure rate for NGS assays attempted on diagnostic prostate biopsies, the most common source of material for genomic testing in routine clinical practice, either due to poor quantity or quality of tumor DNA.

Functional tissue-based assays are emerging as promising biomarkers to assess HRR functionality in clinical samples. RAD51 is a protein involved in the final steps of repairing double-strand DNA breaks through homologous recombination. RAD51 forms filaments on single-stranded DNA regions generated during the repair process, facilitating the search for homology chains and strand pairing. Upon double-strand DNA damage, RAD51 foci are detectable in the nucleus, making it a reliable marker for assessing the functionality of HRR in preclinical models. In organoids and patient-derived xenografts, lack of RAD51 foci is associated with PARPi sensitivity. However, clinical applicability was limited, as these assays required external induction of DNA damage, either analyzing biopsies post-treatment or requiring *ex vivo* irradiation of the biopsies (Graeser et al., 2010; Mateo et al., 2019; Naipal et al., 2014). Recently, immunofluorescence-based assays have been optimized to allow the study RAD51 foci on clinical formalin-fixed paraffin-embedded (FFPE) tumor samples (Castroviejo-Bermejo et al., 2018; Cruz et al., 2018)(Pellegrino et al., 2022). Functional HRD status defined by lack of RAD51 foci was predictive of platinum response in early TNBC and high-grade ovarian cancer (Compadre et al., 2023; Llop-Guevara et al., 2021). In prostate cancer, we recently reported the predictive value of RAD51-IF to PARPi treatment in a cohort of patients with HRR mutations in the TOPARP-B clinical trial (Carreira et al., 2021). The performance of the assay and prevalence of functional HRD by RAD51-IF in larger, molecularly unselected populations of patients with metastatic prostate cancer is, however, unknown.

Here we present a comprehensive analysis of homologous recombination status by parallel NGS and RAD51-IF evaluation in a molecularly unselected cohort of primary and metastatic biopsies from metastatic prostate cancer patients.

## MATERIALS AND METHODS

### Study design, patients, and samples

Patients were recruited as part of an academic non-interventional molecular characterization study of mPC at the Vall d’Hebron University Hospital, approved by the local IRB (PRAG5248, approval March 2018). All patients provided informed consent. All consecutive patients who met the eligibility criteria (stage IV prostate cancer and fit for systemic therapy) and had at least one tumor tissue sample (archival diagnostic biopsies or newly acquired, imaging-guided, metastatic biopsies) collected until May 2023 and suitable for molecular studies were included in this analysis. Clinical data were captured from electronic patient records and registered into a REDCap database (Harris et al., 2009, 2019).

### Sample processing and DNA extraction

All tissue specimens underwent central review at VHIO Pathology facilities to determine tumor content and appropriateness for NGS and IF, contingent upon the available material. Whenever feasible, sections for both DNA extraction and IF analysis were concurrently obtained from single FFPE blocks. When frozen blocks were available, those were prioritized for WES, and an FFPE block from the same biopsy procedure was used for IF. In cases where the tissue quantity was insufficient for both NGS and IF, targeted NGS was prioritized based on implications for patient care. Saliva and/or blood were collected from all patients to obtain germline DNA. DNA was extracted from FFPE blocks using either the Qiagen AllPrep® DNA/RNA FFPE kit for FFPE-derived samples or using the Maxwell® RSC FFPE Plus DNA Kit (Promega). For frozen blocks, the Qiagen AllPrep DNA/RNA/miRNA Universal Kit was used. Germline DNA was isolated from blood or saliva samples using the QIAamp DNA Mini kit (Qiagen). DNA underwent mechanical fragmentation using a Covaris M220 focused-ultrasonicator, aiming at 150 bp fragment-size, prior to library preparation.

### Targeted next-generation sequencing

Tumor-only capture-based targeted sequencing was performed using the ISO-accredited VHIO-300 targeted panel (Saura et al., 2023). In brief, libraries were prepared using SureSelect XT Human (Agilent) and captured using a customized panel covering exonic regions of 435 genes. Libraries were sequenced in a HiSeq2500 instrument (Illumina), 2×100 paired end. Sequencing reads were aligned against the GRCh37 (hg19) reference genome using BWA (v0.7.17) (Li & Durbin, 2009), and base recalibrated and indel realigned using GATK (v3.7.0) (McKenna et al., 2010) and abra2 (v2.23) (Mose et al., 2019), respectively. For mutations, variant calling was performed with VarScan2 (v2.4.3) (Koboldt et al., 2012) and Mutect2 (Genome Analysis Toolkit (GATK) v4.1.0.0) (McKenna et al., 2010). Frequent single nucleotide polymorphisms (SNPs) were filtered based on the gnomAD database (allele frequency ≤ 0.0001). Only variants identified by both callers, with a minimum of 7 supporting reads, and with a minimum VAF of 5% for SNVs and 10% INDELs were considered. Variant annotation was performed using publicly available databases (COSMIC, ClinVar, VarSome, OncoKB) and manually curated. Copy number alterations (CNA) were calculated using CNVkit (v0.9.6) (Talevich et al., 2016); categorical annotation of CNA followed the approach reported by Grasso et al (Grasso et al., 2012)..

### Whole-exome sequencing

Libraries were generated using the KAPA HyperPrep kit (Roche) following manufacturer’s instructions and captured with KAPA HyperExome following manufacturer’s instructions (KAPA HyperCap workflow v3, Roche). Sequencing of paired tumor and normal libraries was performed on Illumina HiSeqX or NovaSeq6000 (Illumina) with 150 bp paired-end reads. Reads were mapped to the human reference genome (GRCh38) using the BWA-MEM algorithm (v0.7.15) (Li & Durbin, 2009). BAM files were generated, all duplicate reads were marked, and the quality scores were recalibrated by using Picard (v2.26.2, https://broadinstitute.github.io/picard) and Genome Analysis Toolkit’s Table Recalibration tool (v4.2.5.0) (McKenna et al., 2010), respectively. Somatic mutations were called (tumor versus matched normal) using Mutect2 (Genome Analysis Toolkit (GATK) v4.2.5.0) (McKenna et al., 2010), Strelka2 (v2.9.2) (Kim et al., 2018), and Varscan2 (v2.4.3) (Koboldt et al., 2012) and left-aligned and normalized using bcftools (v1.17) (Danecek et al., 2021a). Mutations detected by at least two algorithms were retained. Annotation of variants was performed using Cancer Genome Interpreter (CGI, https://www.cancergenomeinterpreter.org/analysis) followed by manual curation. Germline mutations were called with Varscan (v2.4.3) and annotated with Annovar (v2.27.1) (Wang et al., 2010). Allele-specific copy-number profiles were estimated from WES data by using ASCAT (v3.1.2) (Van Loo et al., 2010). Low-Pass WGS (LP-WGS, 0.5x) was used for validation of copy-number alterations from both targeted or whole-exome sequencing data in a subset of 53 genes of relevance for prostate cancer (Supp Table 1). Reads were mapped to the hg19 human genome with Bowtie2 (v2.3.5.1) (Langmead et al., 2009); duplicates were removed using SAMtools (v1.10) (Danecek et al., 2021b); segmentation was performed with ReadCounter (HMM copy utils R package) (Lai et al., 2023) with a 500Kb window, removing low-quality reads (<Q20); ichorCNA was used to calculate segment-based copy number (Adalsteinsson et al., 2017). Homozygous deletion (HOMDEL) was defined as estimation of 0 copies, heterozygous deletion (HETDEL) as 1 copy, gain between 3 and 6 copies, and amplification (AMP) > 6 copies.

**Table 1.** Disease characteristics at the time of diagnosis.

| Table 1. Disease characteristics at the time of diagnosis |  |
| --- | --- |
| Characteristic | Value |
| n | 187 |
| Histology, n (%) |  |
| Adenocarcinoma (acinar) | 180 (96.3) |
| Other | 7 (3.7) |
| Histologic Grade Group, n (%) |  |
| 1-3 | 37 (19.7) |
| 4 | 56 (29.9) |
| 5 | 77 (41.2) |
| Unknown | 17 (9.1) |
| PSA (ng/ml), median [IQR] | 38.89 [12.28, 264.75] |
| Tumor category, n (%) |  |
| T1 | 5 (2.7) |
| T2 | 18 (9.7) |
| T3 | 75 (40.5) |
| T4 | 41 (22.2) |
| Tx | 46 (24.9) |
| Nodal stage, n (%) |  |
| N0 | 51 (27.4) |
| N1 | 94 (50.5) |
| Nx | 41 (22.0) |
| Metastases, n (%) |  |
| M0 | 68 (36.4) |
| M1 | 117 (62.6) |
| Mx | 2 (1.1) |
| Stage, n (%) |  |
| I-II | 11 (5.8) |
| IIIA | 4 (2.1) |
| IIIB | 23 (12.3) |
| IIIC | 9 (4.8) |
| IVA | 19 (10.2) |
| IVB | 117 (62.6) |
| Unknown | 4 (2.1) |
| REFERENCES. PSA: Prostate-specific antigen, IQR: interquartile range. |  |

### Genomic scars – HRD scores

Loss of heterozygosity (HRD-LOH), large-scale state transitions (LST), number of telomeric allelic imbalances (NtAI) and the unweighted numeric sum of LOH, tAI, and LST, named HRD-sum, were determined from WES using the scarHRD R package (Sztupinszki et al., 2018). Additionally, these three genomic scars were also determined from the capture-based NGS panel sequencing data adapting the algorithm previously described by Marquard et al (Marquard et al., 2015) using the B-allele fraction calculation determined with the 10^5^ SNPs distributed throughout the genome from the backbone of the VHIO-CARD-300 panel. From LP-WGS, large-scale genomic alteration (LGA) was estimated with shallowHRD (Eeckhoutte et al., 2020).

### RAD51 Immunofluorescence

IF for RAD51, geminin (GMN), and phospho-histone H2AX (γH2AX) was performed on 3 μm FFPE tumor biopsy sections following previously reported methods (Castroviejo-Bermejo et al., 2018; Cruz et al., 2018). γH2AX was used as quality check of DNA double-strand break in the tumor; RAD51 foci were quantified only in GMN positive cells, that would correspond to those cells in the S–G2 cell-cycle phase, when HRR takes place.

For target antigen retrieval, sections were microwaved in DAKO Antigen Retrieval Buffer pH 9.0. Sections were permeabilized with DAKO Wash Buffer for 5 minutes, followed by five-minute incubation in blocking buffer (DAKO Wash Buffer with 1% BSA). Primary antibodies were diluted in DAKO Antibody Diluent and incubated at room temperature for 1 hour. Sections were washed and blocked again. Secondary antibodies were diluted in blocking buffer and incubated for 30 minutes at room temperature. Finally, sections were dehydrated with increasing concentrations of ethanol and mounted with DAPI ProLong Gold Antifade Reagent (Invitrogen). The following primary antibodies were used: rabbit anti-RAD51 (Abcam ab133534, 1:1,000), mouse anti-GMN (NovoCastra NCL-L, 1:60), rabbit anti-GMN (ProteinTech 10802–1-AP, 1:400), and mouse anti-γH2AX (Millipore #05636, 1:200) and mouse anti-BRCA1 (Santa Cruz Biotechnology Inc, Dallas, TX sc-6954, 1:50). Goat anti-rabbit Alexa Fluor 568 (Invitrogen; 1:500), goat anti-mouse Alexa Fluor 488 (Invitrogen; 1:500), donkey anti-mouse Alexa Fluor 568 (Invitrogen; 1:500), and goat anti-rabbit Alexa Fluor 488 (Invitrogen; 1:500) were used as secondary antibodies.

Scores were assessed on life images using a 60× immersion oil objective with a Nikon Eclipse Ti-E microscope. To be considered evaluable, specimens should present at least 40 GMN-positive cells, and more than 25% of γH2AX/GMN–positive tumor cells. RAD51 was quantified in tumor areas by scoring the percentage of GMN-positive cells with five or more RAD51 nuclear foci. Scoring was performed blinded to the genomics data. IF images were acquired with a 60× objective using a Nikon DS-Qi2 digital camera and generated using NIS-Elements-AR (version 4.40) software.

### Statistical analysis

Frequencies and distributions are reported as descriptive statistics. Comparisons for continuous response variables were performed by t-test or Wilcoxon test according to the test assumptions being fulfilled; besides, linear models were used, particularly, when covariables adjustment was required. To test the association between categorical variables, the Fisher’s Exact Test was applied. Logistic regression was implemented to analyze binary response variables building simple and multiple models. In this setting, results were presented along with the odds ratio (OR) and the 95% confidence interval (95% CI). Correlations were assessed by Pearson correlation coefficient.

For analyses including RAD51 IF data, samples were classified as RAD51 “high” or “low” by applying a previously reported cutoff of 10% RAD51-positive/geminin-positive tumor cells (Castroviejo-Bermejo et al., 2018; Cruz et al., 2018). To explore and define the variables associated with a response variable, such as the set of genes linked with the RAD51 assay result, two approaches were considered. First, a variable selection method followed by the analysis of the potential variables in a multiple model. To complete this approach, lasso was run defining the lambda parameter by leave-one-out cross validation and considering both the minimum value and the minimum plus 1 standard deviation. The selected variables were included in a multiple regression logistic model to finally test their significance and to determine the OR and the 95% CI. Secondly, as an additional approach, a random forests model was built including all the variables to obtain variable importance metrics. To build all these models, a fixed set of genes of interest (n=53) were included as binary predictors as follows: tumor suppressor genes were considered altered when loss of function alterations were detected (such as deep deletions and/or mutations, but not amplifications); oncogenes were considered altered when amplifications and/or mutations were identified (not deletions); and for other genes, such as those with chromatin remodeling function, all alterations considered pathogenic were included.

No imputation was done in any step. The significance level was alpha=0.05 two-sided for all tests. The statistical analysis was performed with R, version 4.3.0 (R Core Team, 2022). Libraries used are listed in the references (Allaire et al., 2023; Breiman et al., 2022; Chen, 2022; Friedman et al., 2010, 2023; Garnier et al., 2023; Garnier, 2023; Gu, 2023; Gu et al., 2016; Kennedy, 2020; Liaw & Wiener, 2002; Neuwirth, 2022; Pedersen, 2022; Prabhakaran, 2016; R Core Team, 2022; Simon et al., 2011; Tay et al., 2023; Wickham, 2023; Wickham et al., 2019; Xie, 2014, 2015, 2023; Xie et al., 2018, 2020; Yoshida & Bartel, 2022).

## RESULTS

### Study population and sample disposition

The study included 219 samples from 187 advanced prostate cancer patients. At diagnosis, most patients presented tumors with high Gleason grade (group 4-5 tumors, n=133, 71.1%) and *de novo* metastatic disease (n=117, 62.6%) (Table1). The samples were acquired from primary (n=151, 68.9%) and metastatic tumor lesions (n=68, 31.1%) (Table 2). Metastatic samples were acquired mostly from bone (n=33, 15.1% overall study samples) and lymph nodes (n=23, 10.5%). In 169 cases (77.2%), the biopsy had been collected prior to hormonal therapy (hormone-sensitive, HSPC), whereas in 50 cases (22.8%), the biopsy was acquired upon castration-resistance (CRPC).

**Table 2.** Tumor and Clinical characteristics at the time of biospecimen acquisition.

|  |  |
| --- | --- |
| Characteristic | Value |
| n | 219 |
| Biopsy site, n (%) |  |
| Prostate | 148 (67.6) |
| Bone | 33 (15.1) |
| Lymph node | 23 (10.5) |
| Liver | 9 (4.1) |
| Urinary bladder | 2 (0.9) |
| Lung | 1 (0.5) |
| Penis | 1 (0.5) |
| Pleura | 1 (0.5) |
| Urethra | 1 (0.5) |
| Tumor site category, n (%) |  |
| Metastasis | 68 (31.1) |
| Primary tumor | 151 (68.9) |
| Hormone sensitivity status, n (%) |  |
| CRPC | 50 (22.8) |
| HSPC | 169 (77.2) |
| Clinical setting (%) |  |
| Diagnosis | 144 (65.8) |
| Relapse after local treatment | 8 (3.7) |
| Adv disease, on first hormone naïve therapy | 18 (8.2) |
| Adv disease, castration resistant- 1st line | 12 (5.5) |
| Adv disease, castration resistant- 2nd or subsequent lines | 37 (16.9) |
| REFERENCES. CRPC: castration resistant prostate cancer, HSPC: hormone sensitive prostate cancer, Adv disease: advanced disease. |  |

All samples (n=219) underwent NGS: targeted sequencing was performed in 156 biopsies from 151 patients and WES in 87 biopsies from 71 patients; 38 biopsies underwent both targeted and whole-exome sequencing (Fig.1A). Genomics data was obtained for 181 samples (157 patients): 139 samples (134 patients) had targeted sequencing results, whereas WES data was successfully obtained for 80 samples (65 patients) (Fig.1B).

**Figure 1.**
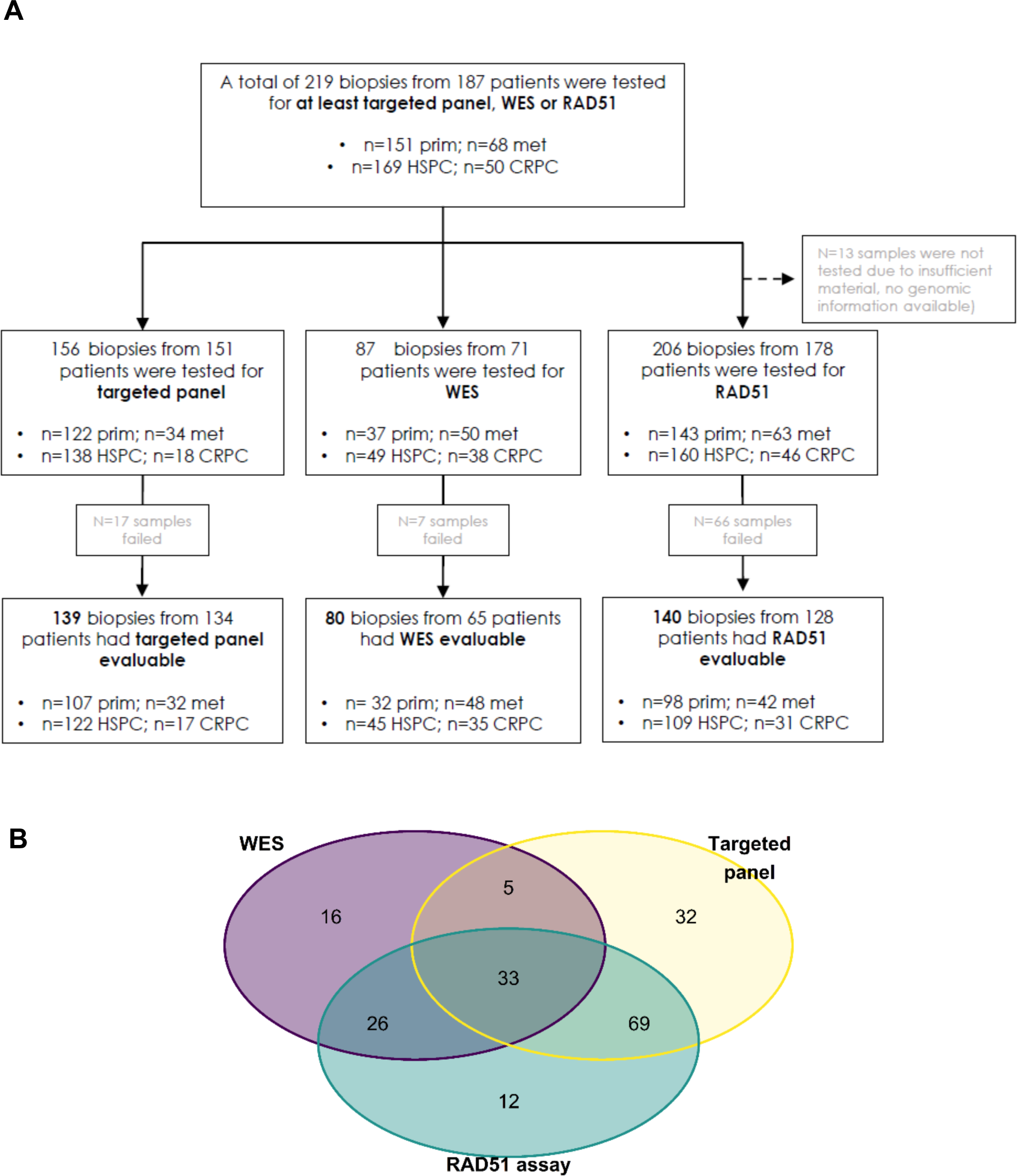
**Sample disposition in this study per assay. A**, Diagram of the “intention-to-test” population and overview of sample disposition. Prim: primary tumor; met: metastatic tumor; HSPC: hormone-sensitive prostate cancer; CRPC: castration-resistant prostate cancer; pt: patient; WES: whole exome sequencing. **B**, Venn diagram for evaluable samples for genomic and/or functional assays (WES n=80, targeted panel n=139, RAD51 assay n=140).

Out of 219 samples, RAD51 immunofluorescence was performed on 206 biopsies from 178 patients, as for the remaining 13 samples there was insufficient material for IF after NGS. The test was informative in 68% of the samples (140/206), namely for 98 primary tumors and 42 metastatic tumor biopsies. Reasons for non-evaluability included insufficient tumoral cells, low proliferative tumors (insufficient geminin-positive cells) or low levels of DNA damage (γH2AX positive cells).

### Genomic characterization of the study population

We investigated the presence of pathogenic mutations and copy number changes on relevant genes involved in the AR pathway, DDR (including HRR, MMR, and others), cell cycle regulation, PI3K and Wnt pathways, among others (Fig. 2). The most frequently altered genes included *TP53* (40%), *AR* (15%), *PTEN* (14%), *FOXA1* (12%), *MYC* (10%) and *BRCA2* (9%). As expected, *AR* amplifications were observed primarily in samples collected at the castration-resistant setting (37%) compared to HSPC (4%; p-value<0.0001). Detailed genomic information by targeted vs whole-exome sequencing is shown in Supp Fig. 1 A-B.

**Figure 2.**
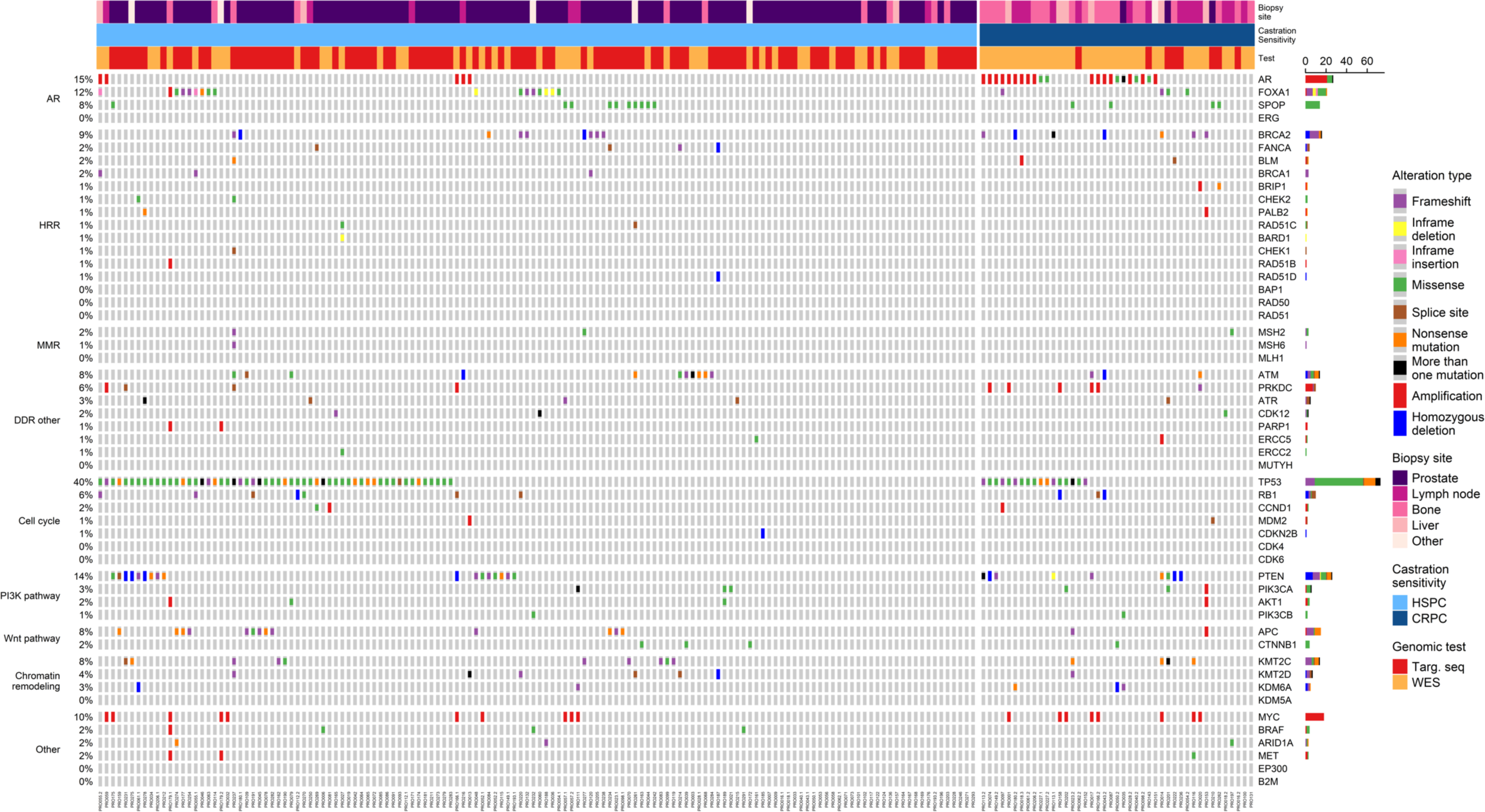
The landscape of genomic alterations in study cohort (n=181). Oncoprint of pathogenic mutations (SNVs and Indels) categorized by coding consequence and copy changes (amplifications and homozygous deletions) across the entire cohort grouped by castration sensitivity. HSPC: hormone-sensitive prostate cancer; CRPC: castration-resistant prostate cancer.

Regarding alterations in HRR-related genes, the most frequently altered was *BRCA2* (9%, including frameshift mutations and homozygous deletions), followed by *BRCA1* and *FANCA* (2% each). Among other DDR genes, *ATM* was the most altered gene (8%). The prevalence of DDR mutations was similar among HSPC and CRPC samples.

### Homologous recombination repair function based on RAD51 IF

The prevalence of tumor cells with RAD51 nuclear foci was quantified in 140 biopsies from 128 patients (Fig. 3A). The median RAD51 IF score was 28.5 (IQR 13.9 - 43.3). Applying a previously defined threshold of ≤10% RAD51-positive cells to be considered HRR deficient, 21% of the samples (30/140) were classified as HRD or “RAD51 low”. No differences were observed when comparing primary (n=98) versus metastatic (n=42) tumors (median 29.6 and 27.0 respectively; p-value=0.704), nor hormone-sensitive (n=109) versus castration-resistance (n=31) (29.0 and 28.0 respectively; p-value=0.49; Fig. 3B).

**Figure 3:**
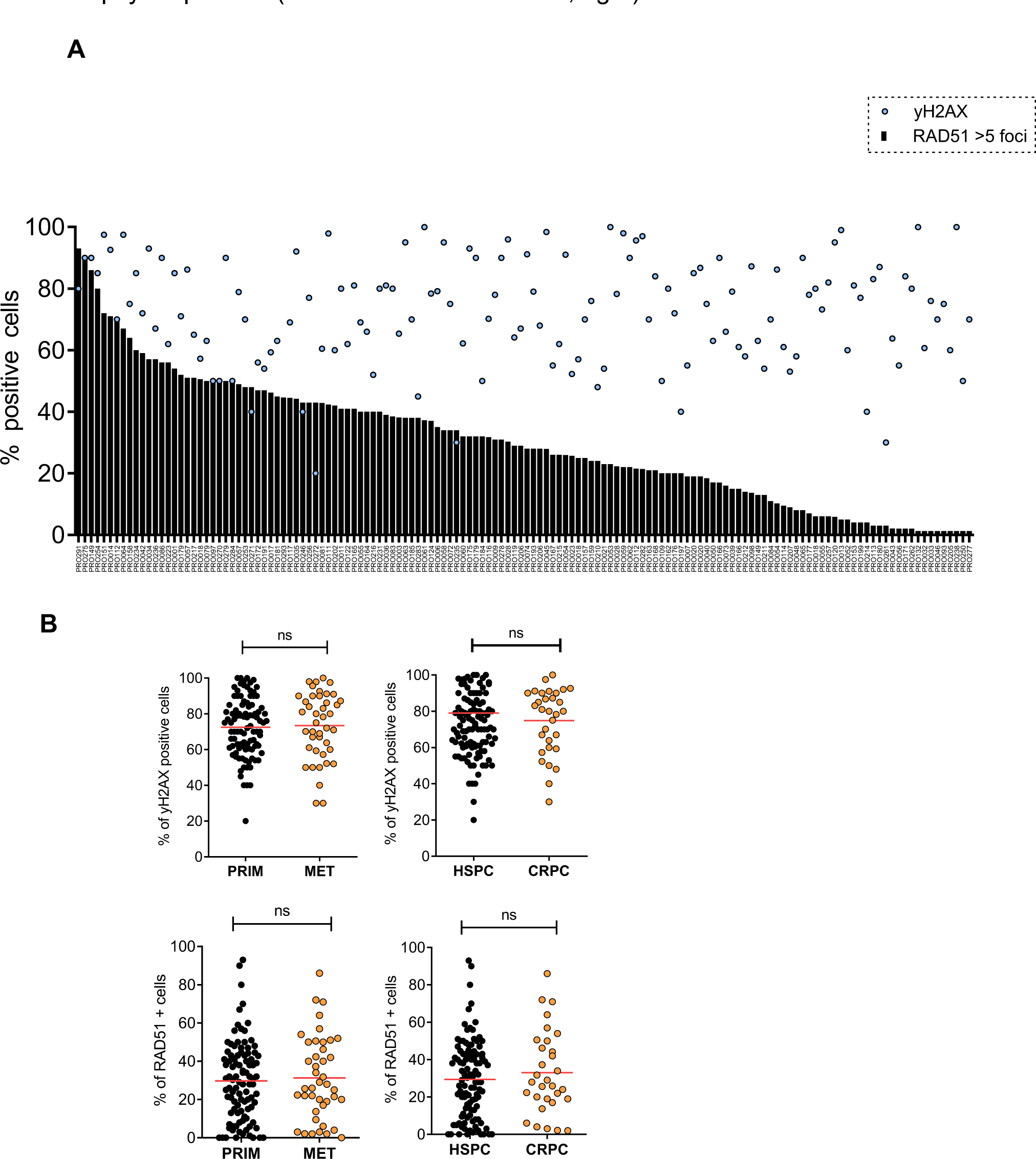
Prevalence of RAD51 scores on the patient cohort. A, Barplot depicting RAD51 foci (bars) and γH2AX (dots) distribution across the evaluable samples. B, Comparison of percentages of γH2AX positive cells (top panels) and RAD51 positive cells (lower panels) by IF in primary vs metastatic biopsies (left) or based on the hormonal therapy exposure of the patient at the time of biopsy acquisition (hormone naive vs CRPC, right).

### Low RAD51 IF scores associate with HRR gene alterations

Figure 4 shows the genomic landscape among the 128 samples with evaluable RAD51 IF, sorted by RAD51 IF score. *TP53* was altered in 44% of the cases followed by *PTEN* (16%), *AR* (15%), *MYC* (14%), *FOXA1* (12%), *BRCA2* (9%), and *ATM* (9%), as the more common alterations.

**Figure 4.**
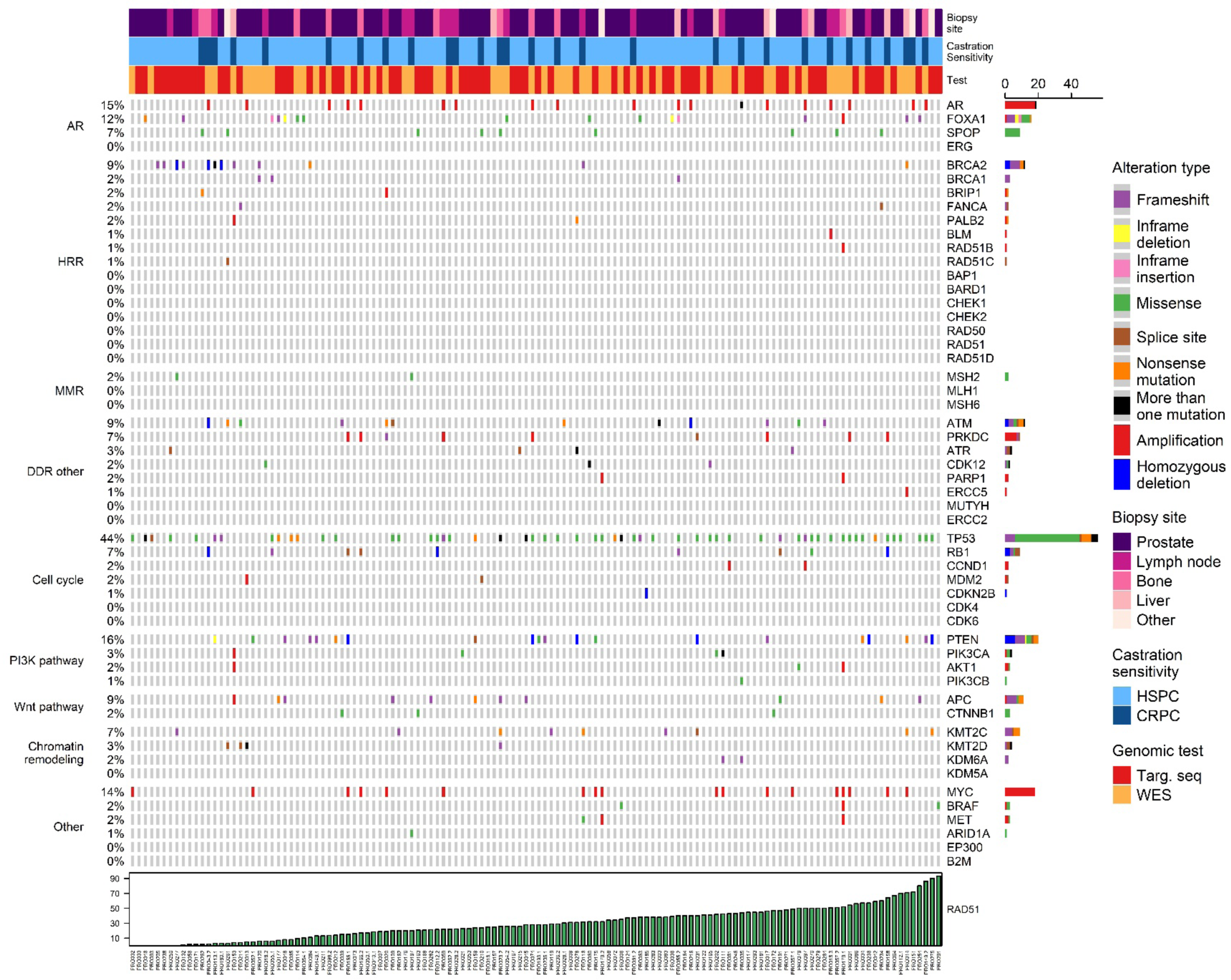
Correlation between NGS and RAD51-IF scores. Oncoprint of pathogenic mutations (SNVs and Indels) categorized by coding consequence and copy changes (amplifications and homozygous deletions), sorted by RAD51 scores from 0 (left) to 100% (right) positive cells in the sample, and color coded by variant category. HSPC: hormone-sensitive prostate cancer; CRPC: castration-resistant prostate cancer.

Cases with pathogenic BRCA1/2 alterations associated with lower RAD51 IF scores, with a median score of 3.5 (IQR 9.8 – 8.5) for *BRCA1/2* altered (n=14) and 29.7 (IQR 19.0 - 44.5) for *BRCA1/2*-WT (n=114) (Figure 5). Considering the threshold of 10% RAD51 positive cells, the sensitivity and specificity of the RAD51 IF assay in identifying *BRCA1/2*-altered cases was 0.71 and 0.85 respectively. When including other HRR genes (*BRIP1, FANCA, PALB2, BLM, CHEK2, RAD50*), the sensitivity and specificity of the RAD51 IF assay were 0.68 and 0.87, respectively (Table 3).

**Figure 5.**
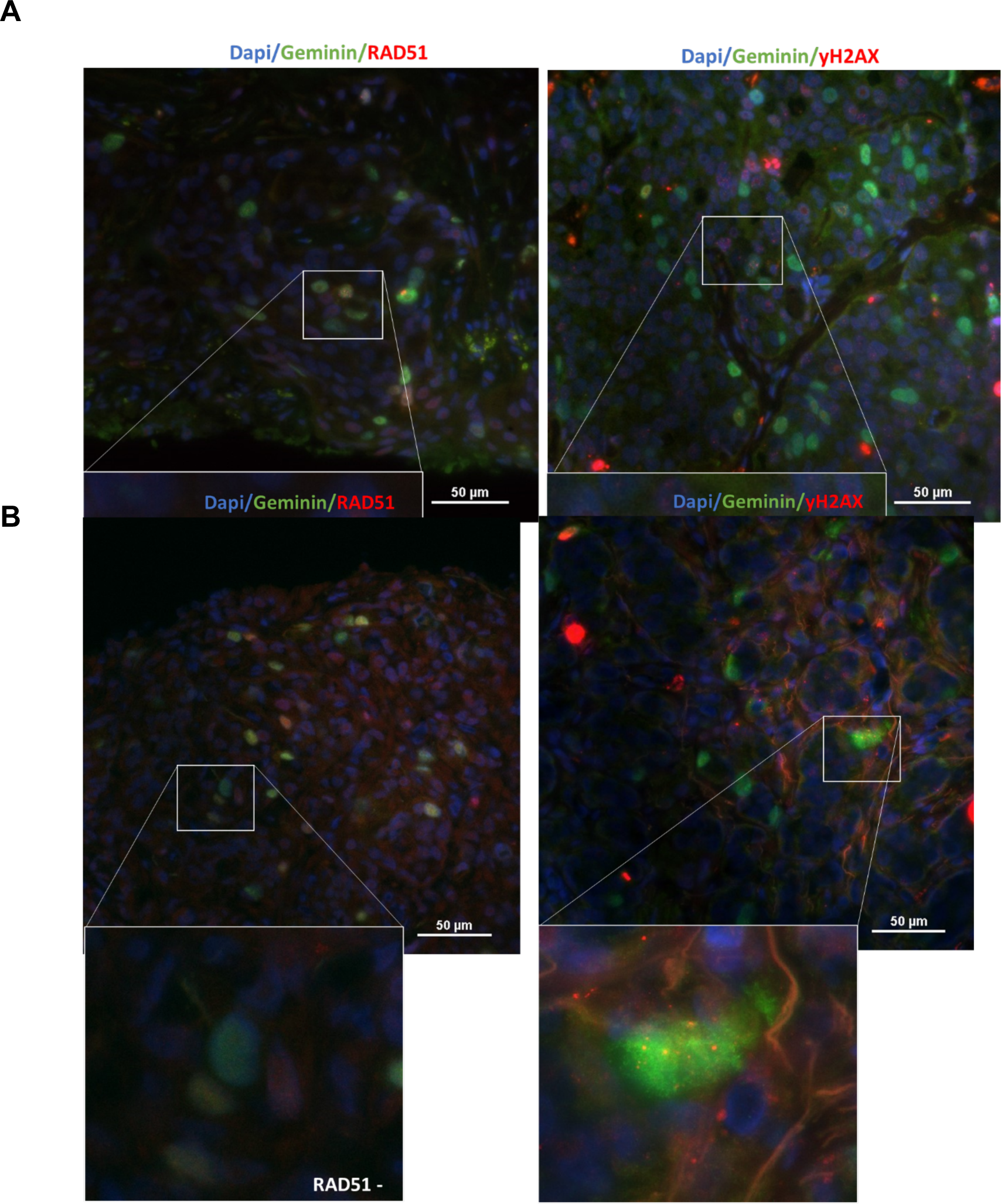
Representative cases correlating RAD51-IF score and genomic characterization. Images of the RAD51 and yH2AX staining from **A**, a RAD51 high sample with no *BRCA1/2* alterations and **B**, a RAD51 low sample with a *BRCA2* alteration.

**Table 3.** Relationship between RAD51 assay result and genomics.

| Table 3. Relationship between RAD51 assay result and genomics. |  |  |  |  |
| --- | --- | --- | --- | --- |
| Confusion matrix |  |  |  |  |
|  | *BRCA1/2* altered | *BRCA1/2* WT | HRR altered | HRR WT |
| RAD51 ≤10 | 10 | 17 | 13 | 14 |
| RAD51 >10 | 4 | 97 | 6 | 95 |
| Performance metrics |  |  |  |  |
| Sensitivity | 71.4 |  | 68.4 |  |
| Specificity | 85.1 |  | 87.2 |  |
| Accuracy | 83.6 |  | 84.4 |  |
| REFERENCES. *BRCA1/2*: *BRCA1* and *BRCA2*, WT: wild type, HRR: HRR genes (*BRCA1*, *BRCA2*, *BAP1*, *BARD1*, *BLM*, *BRIP1*, *CHEK1*, *CHEK2*, *FANCA*, *PALB2*, *RAD50*, *RAD51*, *RAD51B*, *RAD51C*, *RAD51D*). |  |  |  |  |

Delving into the alteration type of these fourteen *BRCA1/2-*altered cases, 3/3 cases with *BRCA2* deep deletion were identified as HRD with the RAD51-IF test. Out of eleven cases with pathogenic BRCA1/2 mutations (frameshift or nonsense), 7 had a RAD51-IF status of HRD and one sample had a score of 11, close to the threshold. The remaining three BRCA1/2 altered cases presented high RAD51-IF values of 31, 40 and 71 respectively (Supp Table 2). We hypothesized that these cases with discrepant results may represent clinical settings where a functional test may add valuable information to NGS. In one of these patients (PRO014), as example, a pathogenic *BRCA2* alteration (BRCA2 Y3006*) was detected, and germline origin was confirmed in subsequent testing (Fig 6A). Based on this finding, the patient was treated with carboplatin, after having progressed to ADT, abiraterone, radium223, docetaxel, and cabazitaxel. The patient presented a PSA and radiographic response to carboplatin, followed by radiographic progression after 7.4 months of treatment. A liver metastatic biopsy was obtained at progression, and the RAD51 IF score was 71%, suggesting the assay might be able to capture HRR dynamics and secondary HRR function restoration. While WES of the liver biopsy did not identify reversion mutations, arguably because of limited coverage in the region, a cfDNA sample collected contemporaneously to the biopsy was sequenced with an external NGS panel, identifying two *BRCA2* reversion mutations (*BRCA2* Y3006Y VAF 0.13, *BRCA2* Y3006L VAF 0.04), in addition to the germline mutation (*BRCA2* Y3006* VAF 0.62) and in line with the secondary resistance to platinum-based chemotherapy.

**Figure 6.**
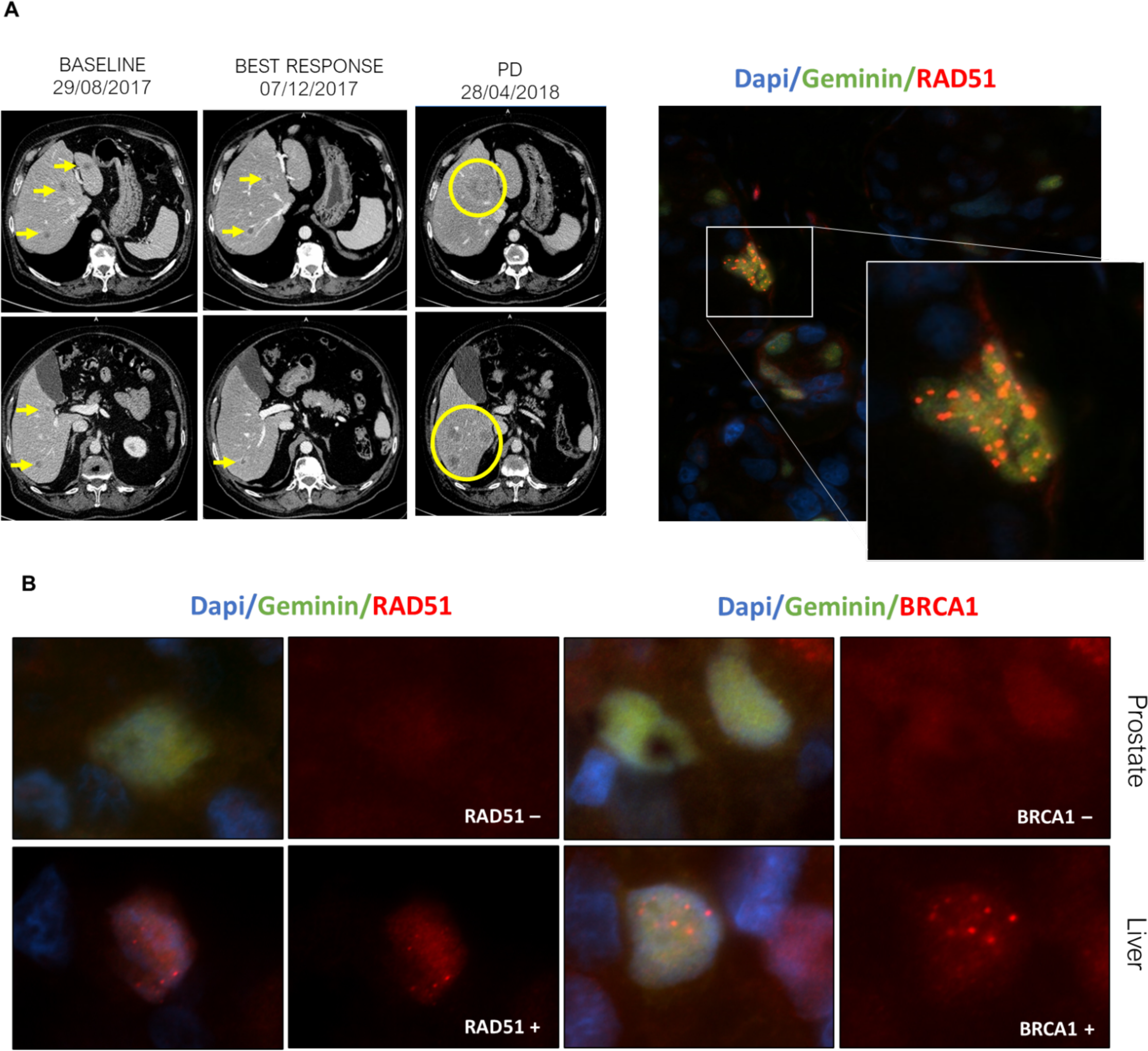
**RAD51 immunofluorescence captures intra-patient heterogeneity. A**, Liver biopsy of a patient with a BRCA2 mutation after progression to plati num-based chemotherapy that shows high percentage of cells positive for RAD51 foci; contemporaneous ctDNA analysis demonstrated *BRCA2* reversion mutations. CT scan (left) of the liver lesions of the patient from baseline, response, and progression to carboplatin. Liver lesions are highlighted in yellow. Representative image of the RAD51 positivity (right) by IF. **B**, Prostate and liver biopsies of a patient with a somatic *BRCA1* mutation detected by NGS. The primary prostate tumor shows RAD51 negative cells but the liver metastasis shows high RAD51 score, in parallel to BRCA1 expression by IF in this liver lesion, but not in the prostate tumor, suggesting restore of BRCA1 expression in the metastases.

Another case (PRO055) where the RAD51-IF assay highlighted a relevant tumor feature was a patient who harbored a *BRCA1* somatic mutation without evidence of second-allele loss in both the primary and metastatic tumors (collected before and after treatment with ADT) and presented a low RAD51 score (6%) on the primary mHNPC sample but a high RAD51 score (40%) in the liver mCRPC biopsy. (Fig. 6B). BRCA1 nuclear foci formation by IF confirmed BRCA1 expression in the liver metastases but not in the primary tumor, arguably explaining the different HRR functional states.

In general, however, cases with matched primary-metastatic samples showed rather consistent RAD51-IF status. Of a total of 10 patients with two or more samples assessed for RAD51 IF and NGS data (Supp Table 1), 8/10 patient-matched pairs of samples were equally classified regarding their functional HRR capacity.

A lasso regression was performed to identify genes associated to RAD51-IF status followed by a multiple regression analysis. All RAD51-evaluable samples were included (n=128), with a predefined set of 53 genes of interest for prostate cancer (Supp Fig. 2). Out of 53 genes, 8 were initially selected by lasso regression: *AR, FOXA1, KMT2C, KMT2D, BRCA1, BRCA2, BRIP1,* and *MET*. Among them, 4 genes were found to significantly associate with RAD51-low status. Three genes increased the odds of RAD51-low result: *BRCA2* (*p*-value<0.001, OR 77.05), *KMT2D* (*p*- value=0.012, OR 103.73), and *FOXA1* (*p*-value=0.033, OR 3.87); whereas alterations in *KMT2C* decreased these odds (*p*-value=0.014, OR 0.02) (Supp Fig. 3). Interestingly, *BRCA2* and *KMT2D* alterations associated with the highest probability (0.91 and 0.93, respectively) of finding a RAD51 low result. Additionally, we orthogonally validated these results by applying a random forests algorithm. Again, the two most relevant genes for determining RAD51-IF status were *BRCA2* and *KMT2D* (Supp Fig. 4).

Of note, only one patient presented *PALB2* pathogenic mutations, specifically a case with a nonsense mutation in *PALB2* with a VAF of 0.20 in a sample with a RAD51-high score. No LOH was detected in the region, suggesting a retained WT allele, that would be sufficient for HRR function, in line with findings in our previous study (Carreira et al., 2021).

### Integration of genomic scars, RAD51 IF, and BRCA1/2 mutational status

We studied the distribution of genomics scars derived from WES (LOH, LST, NtAI, and HRD-sum; n=80), LP-WGS (LGA; n=133), and targeted panel (HRD-sum; n=134) data (Supp Fig. 5). Asymmetric distributions were observed for all variables, in line with previous reports (Supp Fig. 6). The median HRD-sum score was 30.5 (IQR 20.0 - 44.3) from WES and 25.0 (IQR 13.0 – 36.0) from targeted sequencing. A high agreement and correlation were observed between HRD-sum scores from both methods in the subset of 35 samples that underwent both WES and targeted sequencing for cross-validation (Pearson’s coefficient 0.69, *p*-value<0.0001; Supp Fig. 7).

Considering the HRD-sum threshold of ≥42, 27.5% and 20.1% cases were classified as HRD with WES and targeted sequencing, respectively. In line with prior studies by our group and others, we found a statistically significant association for castration sensitivity status with genomic scars: CRPC samples were more likely to be classified as HRD-sum score “high” than HNPC biopsies, both for WES (OR 4.07, *p*- value=0.009) and for targeted panel (OR 5.21, *p*-value=0.003), despite a similar prevalence of HRR gene mutations in these subsets of samples.

The HRD-sum was found significantly associated to *BRCA1/2* mutations (targeted panel HRD-sum: OR 1.05, 95% CI 1.02 - 1.09, *p*-value= 0.004; WES HRD-sum: OR 1.07, 95% CI 1.03 1.12, *p*-value=0.002) (Supp Table 4). When looking at the different components of HRD scar metrics, LST (*p*-value<0.001), NtAI (*p*-value=0.005), and LGA (*p*-value=0.022) were found associated with BRCA1/2 status, but no significant association was observed for LOH (*p*-value=0.28) (Supp Fig. 8-9).

The HRD-sum derived from the targeted panel was found to significantly associate with RAD51-IF status (n=99; categorical classification as low vs high: OR 1.03, 95% CI 1.005 - 1.062; *p*-value=0.021). When looking at HRD-sum based on WES (n=59), there was a non-significant trend for association (OR 10.2, 95% CI 0.988 – 1.061, *p*-value=0.2) that became significant when adjusting for hormone-sensitive vs castration-resistant status (*p*-value=0.035), in line with the described higher HRD scores in castration-resistant prostate cancer.

## DISCUSSION

Molecular stratification has entered prostate cancer care; NGS tests have emerged as a pivotal strategy for patient stratification and treatment selection in metastatic prostate cancer. The mainstay example of clinical utility so far is the association of HRR defects with responses to PARP inhibitors and platinum-based chemotherapy. However, inequalities in access to genomic testing, high NGS technical failure rates for clinical diagnostic prostate biopsies and limited understanding of the functional impact of some of the mutations observed beyond *BRCA1/2* still challenge the deployment of precision medicine in routine mPC care. Functional assays that can detect dynamic changes in HRR capacity have potential to improve patient stratification strategies, and to help prioritize for NGS testing if resources are limited.

In this study, 219 biopsies from 187 mPC patients underwent both functional HRD tissue-based immunofluorescence and NGS assays. Genomic analysis revealed frequent alterations in *TP53, PTEN, AR, MYC*, *BRCA2*, *ATM*, and *BRCA1*, emphasizing the complex genomic landscape of mPC. The RAD51-IF assay was informative in a substantial proportion (68%) of FFPE samples. We observed a clear association between lower RAD51 IF scores and *BRCA2* alterations, suggesting this assay can identify patients with HRR defective prostate cancer; this can be of relevance in cases with limited tissue availability or DNA degradation where NGS is not feasible.

Previous studies suggest that alterations in HRR genes are early events in patients with lethal prostate cancer, being present already in the primary tumors of these patients. In our study, we also found a similar prevalence of HRD phenotype, based on RAD51-IF, when considering primary treatment-naïve or metastatic castration-resistance biopsies. Based on the pre-defined cutoff of ≤10% RAD51-positive tumor cells, 21% of the overall samples were classified as HRD. The IF assay also helped to identify cases where, despite presenting HRR mutations, the tumors may not have developed complete loss of HRR function.

Contrarily the strong association with *BRCA1/2* alterations, there was no association of the RAD51-IF assay with *ATM*-mutated cases, further suggesting that the responses observed to platinum or PARPi in some *ATM*-mutated patients is not necessarily dependent on an HRD synthetic lethal interaction (Neeb et al., 2021).

We also explored genomics scars and their association with RAD51-defined phenotypes. Although genomic scars have demonstrated their potential to guide patient stratification in breast and ovarian cancer, their clinical value is still unclear in prostate cancer. In our study, the association between RAD51 scores and different HRD genomic signatures became stronger when adjusting for castration-resistance status of the biospecimen. This is consistent with prior studies reported by our group and others, showing that treatment resistance and TP53 mutations impact the distribution of genomic scars in prostate cancer, independently to *BRCA1/2* mutations. The distribution of RAD51 IF scores was not impacted by clinical state at the time of sample acquisition. Moreover, in our study, we were able to identify high RAD51 scores upon secondary resistance to carboplatin, arguably supporting the potential of the assay to capture mechanisms of resistance.

These are the first results of the functional assay on a molecularly unselected cohort of mPC patients. Our study cohort was recruited prior to wide PARPi availability in our healthcare system. Hence, the main limitation of the study is the lack of correlation with PARPi clinical responses. However, we previously reported the association between this RAD51 IF assay and response to olaparib in an independent cohort of patients preselected based on HRR gene mutations in the TOPARPB clinical trial (Mateo et al., 2015, 2020)(Carreira et al., 2021).

Overall, this study represents the first implementation of a RAD51-based functional assay in a molecularly unselected cohort of advanced prostate cancer, demonstrating the feasibility in clinical routine samples and its potential to inform patient stratification in metastatic prostate cancer.

## Acknowledgements

This work was funded by an Impact Award from the Department of Defense CDMRP (PC170510P1) and funding by AstraZeneca (ESR-21-21360). The funders had no role in the design or conduction of the study and were not involved in the analysis or interpretation of results.

We also acknowledge support from the CRIS Cancer Foundation (grant TCL_2020- 10 to J. Mateo) and Fundación AECC (LABAE20019MATE). Sara Arce-Gallego was supported by Instituto de Salud Carlos III (FI19/00280). Pablo Cresta was supported by an ESMO Translational Research Fellowship. Alba Llop-Guevara was supported by an AECC grant (INVES20095LLOP). This study was partly funded with an ERA PerMed grant to Violeta Serra (ERAPERMED2019-215). This Research Project was supported by ESMO with the aid of a grant from BMS. Any views, opinions, findings, conclusions, or recommendations expressed in this material are those solely of the author(s) and do not necessarily reflect those of ESMO or BMS. VHIO authors would like to acknowledge: the Spanish State Agency for Research (Agencia Estatal de Investigación) for the financial support as a Center of Excellence Severo Ochoa (CEX2020-001024-S/AEI/10.13039/501100011033), the Cellex Foundation for providing research facilities and equipment and the CERCA Programme from the Generalitat de Catalunya for their support on this research.

We acknowledge Marta Guzman, Olga Rodriguez, Anna Serradell and Maria del Mar Suanes for support conducting experiments and/or collecting data at VHIO.

We would also like to thank all patients who participated in this project by donating their biospecimens and data, for their generous contribution to cancer research.

