## Supplementary material for "Evaluation of homologous recombination repair status in metastatic prostate cancer by next-generation sequencing and functional tissue-based immunofluorescence assays": Suppl. Material

Title:

Table of Content

Supplemental Figure 1. The landscape of genomic alterations according to each genomic test.

Supplemental Figure 2. Alteration frequency for each gene of interest.

Supplemental Figure 3. Forest plot for the multiple model including the four selected genes.

Supplemental Figure 4. Random forests variable importance plot.

Supplemental Figure 5. Venn diagram involving genomic scar results for each test.

Supplemental Figure 6. Frequency of each genomic scar derived from WES, LP-WGS, and targeted panel.

Supplemental Figure 7. Relationship between HRD-sum from WES and targeted panel.

Supplemental Figure 8. Relationship between scars and RAD51.

Supplemental Figure 9. Comparison between RAD51 low and high and each scar.

Supplemental Table 1. Curated list with genes of interest.

Supplemental Table 2. Description of the BRCA1/2 altered cases.

Supplemental Table 3. Cases with more than one sample from the same patient. Descriptive characteristics.

Supplemental Table 4. Regression analysis results for BRCA1/2 as response variable.

Supplemental Table 5. Regression analysis results for RAD51 as response variable.

**Supplemental Figure 1.**

**The landscape of genomic alterations according to each genomic test. A. WES. B. Targeted panel.** Oncoprint of pathogenic mutations (SNVs and Indels) categorized by coding consequence and copy changes (amplifications and homozygous deletions) across the entire cohort grouped by castration sensitivity. HSPC: hormone-sensitive prostate cancer; CRPC: castration-resistant prostate cancer.

# A.

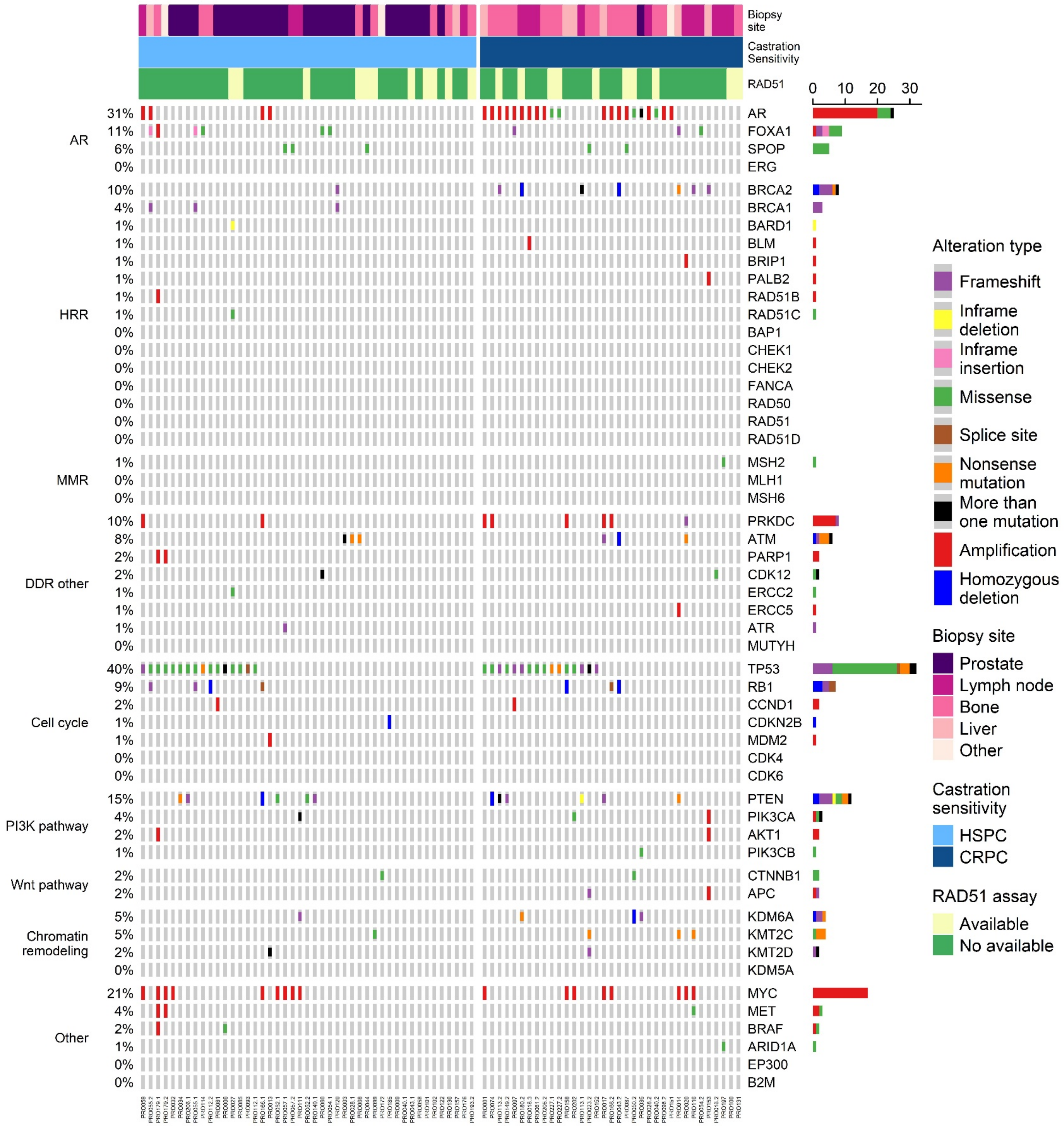

B.

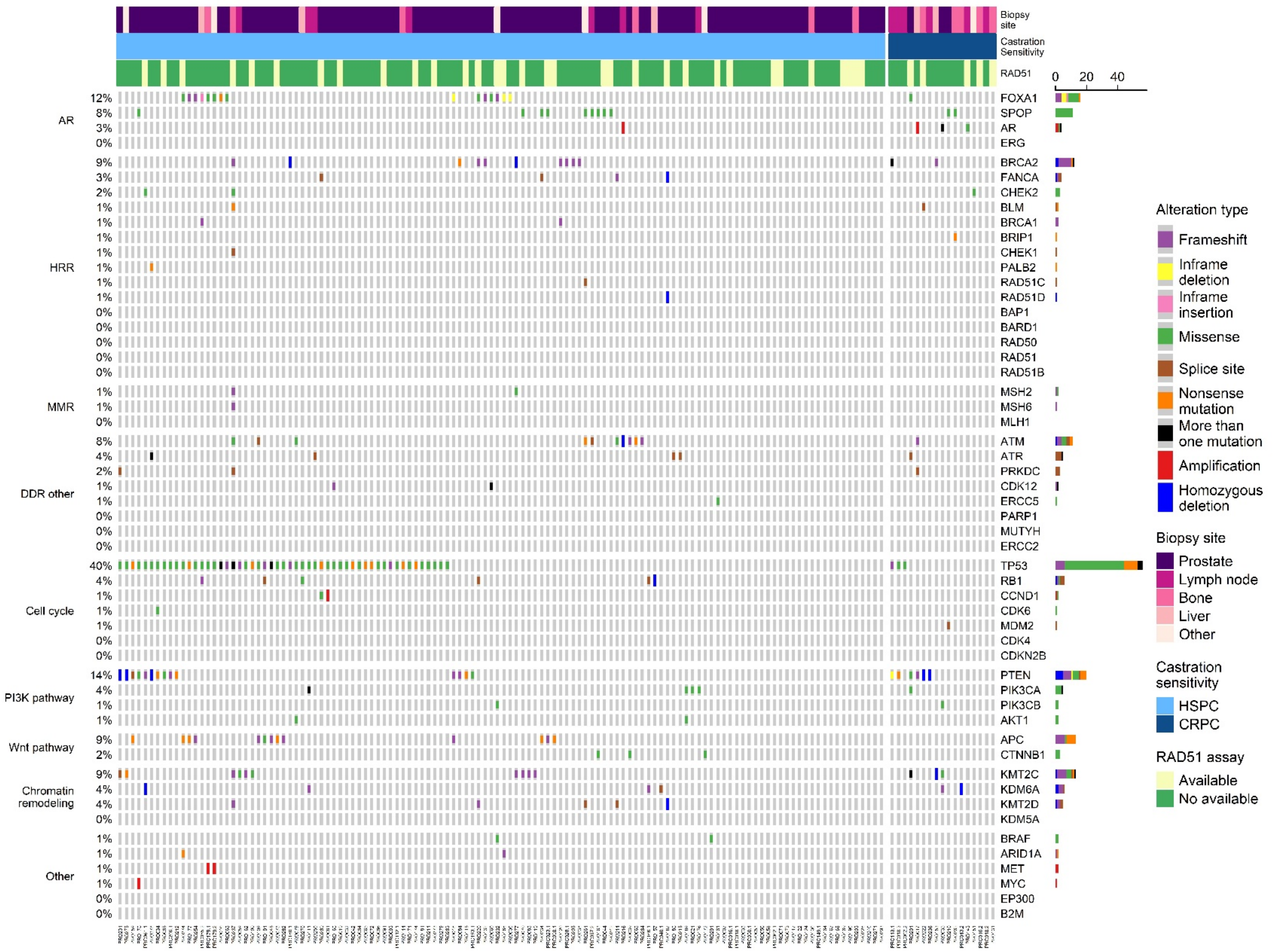

Biopsy site
Castration Sensitivity
RAD51
0 20 40
AR
HRR
MMR
DDR other
Cell cycle
PI3K pathway
Wnt pathway
Chromatin remodeling
Other
FOXA1
SPOP
AR
ERG
BRCA2
FANCA
CHEK2
BLM
BRCA1
BRIP1
CHEK1
PALB2
RAD51C
RAD51D
BAP1
BARD1
RAD50
RAD51
RAD51B
MSH2
MSH6
MLH1
ATM
ATR
PRKDC
CDK12
ERCC5
PARP1
MUTYH
ERCC2
TP53
RB1
CCND1
CDK6
MDM2
CDK4
CDKN2B
PTEN
PIK3CA
PIK3CB
AKT1
APC
CTNNB1
KMT2C
KDM6A
KMT2D
KDM5A
BRAF
ARID1A
MET
MYC
EP300
B2M
Alteration type
Frameshift
Inframe deletion
Inframe insertion
Missense
Splice site
Nonsense mutation
More than one mutation
Amplification
Homozygous deletion
Biopsy site
Prostate
Lymph node
Bone
Liver
Other
Castration sensitivity
HSPC
CRPC
RAD51 assay
Available
No available

**Supplemental Figure 2.**

**Alteration frequency for each gene of interest.** The alteration status was defined according to each gene function. Tumor suppressor genes were considered altered when loss of function alterations were detected (such as deep deletions or mutations but not amplifications); oncogenes, which need a gain of function, were considered altered when amplifications and mutations were identified (not deletions); and for other genes, such as those with chromatin remodeling function, all the pathogenic alterations were considered.

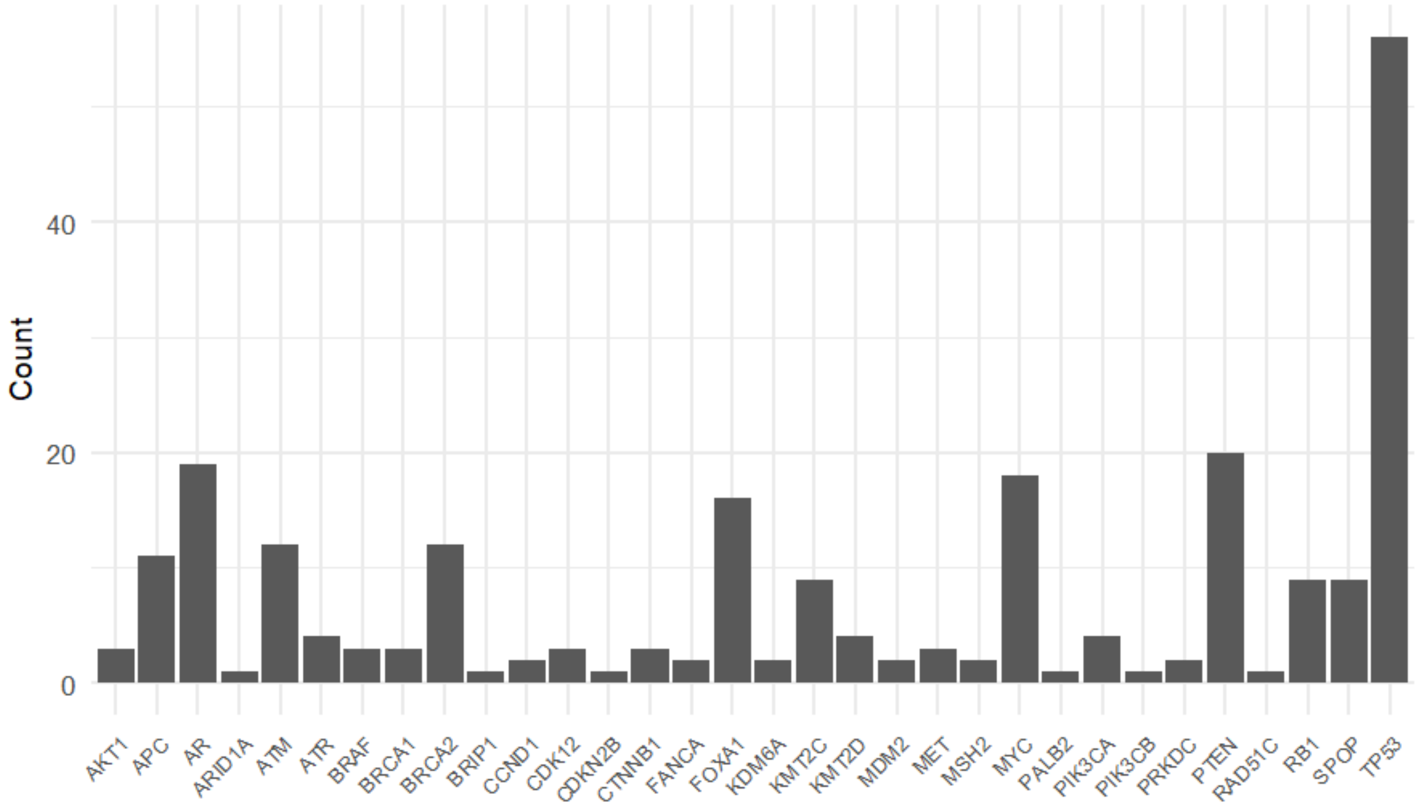

**Supplemental Figure 3.**

**Forest plot for the multiple model including the four selected genes.** KMT2C is associated with a decrease in the odds of obtaining a RAD51 low (OR 0.02). KMT2D and BRCA2 have the highest OR.

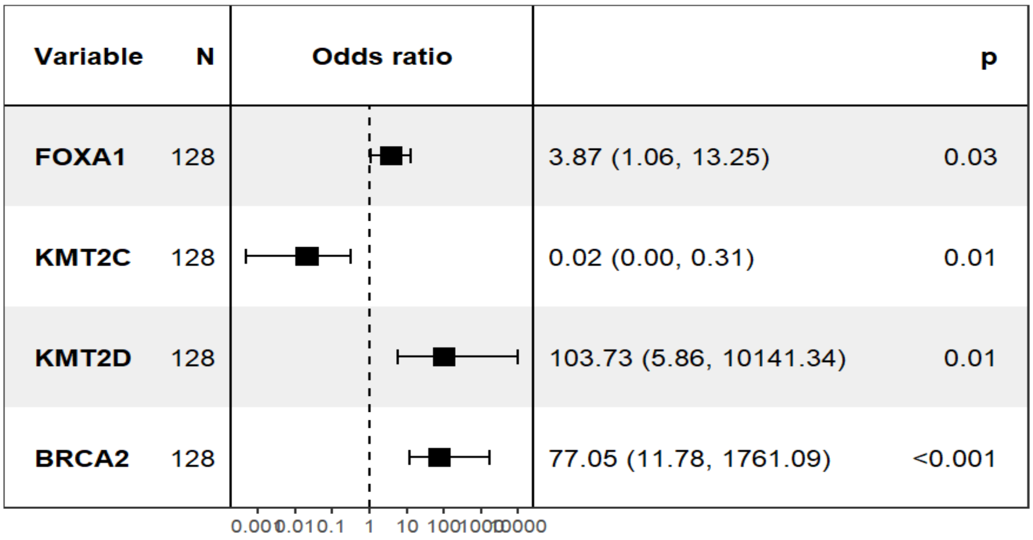

Supplemental Figure 4.

**Random forests variable importance plot.** Genes selected by lasso are highlighted with an asterisk (*) and those included in the final model with two asterisks (**).

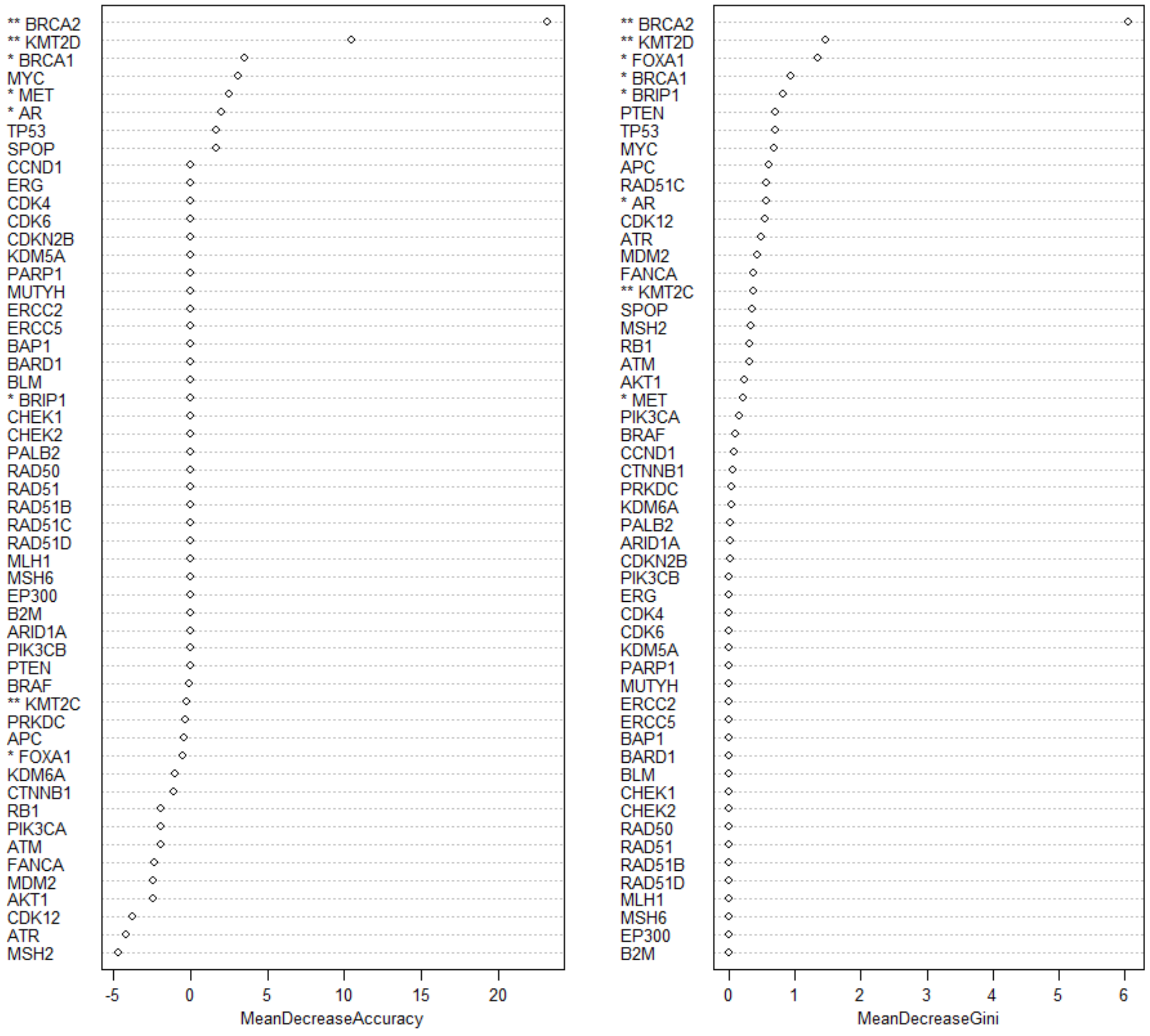

**Supplemental Figure 5.**

**Venn diagram involving genomic scar results for each test (WES, low pass WGS, and targeted panel).** There are in total 80 samples with WES, 134 with targeted panel, and 133 with low pass WGS (LP-WGS).

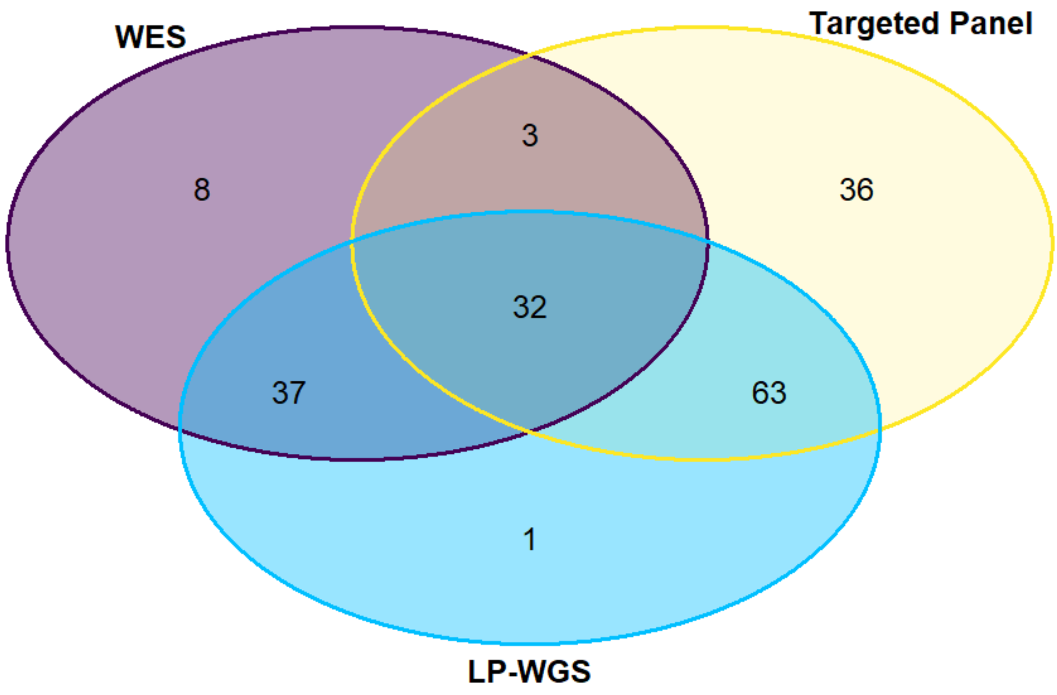

**Supplemental Figure 6.**

**Frequency of each genomic scar derived from WES (violet), LP-WGS (cyan), and targeted panel (yellow). A**. The LOH genomic scar from WES, **B**. The NtAI genomic scar from WES, **C.** The LST genomic scar from WES, **D.** The HRD-sum calculated as the sum of LOH, NtAI, and LST from WES, **E.** The LGA genomic scar from LP-WGS, **F.** The HRD-sum calculated as the sum of LOH, NtAI, and LST from targeted panel. Each plot has not the same scale.

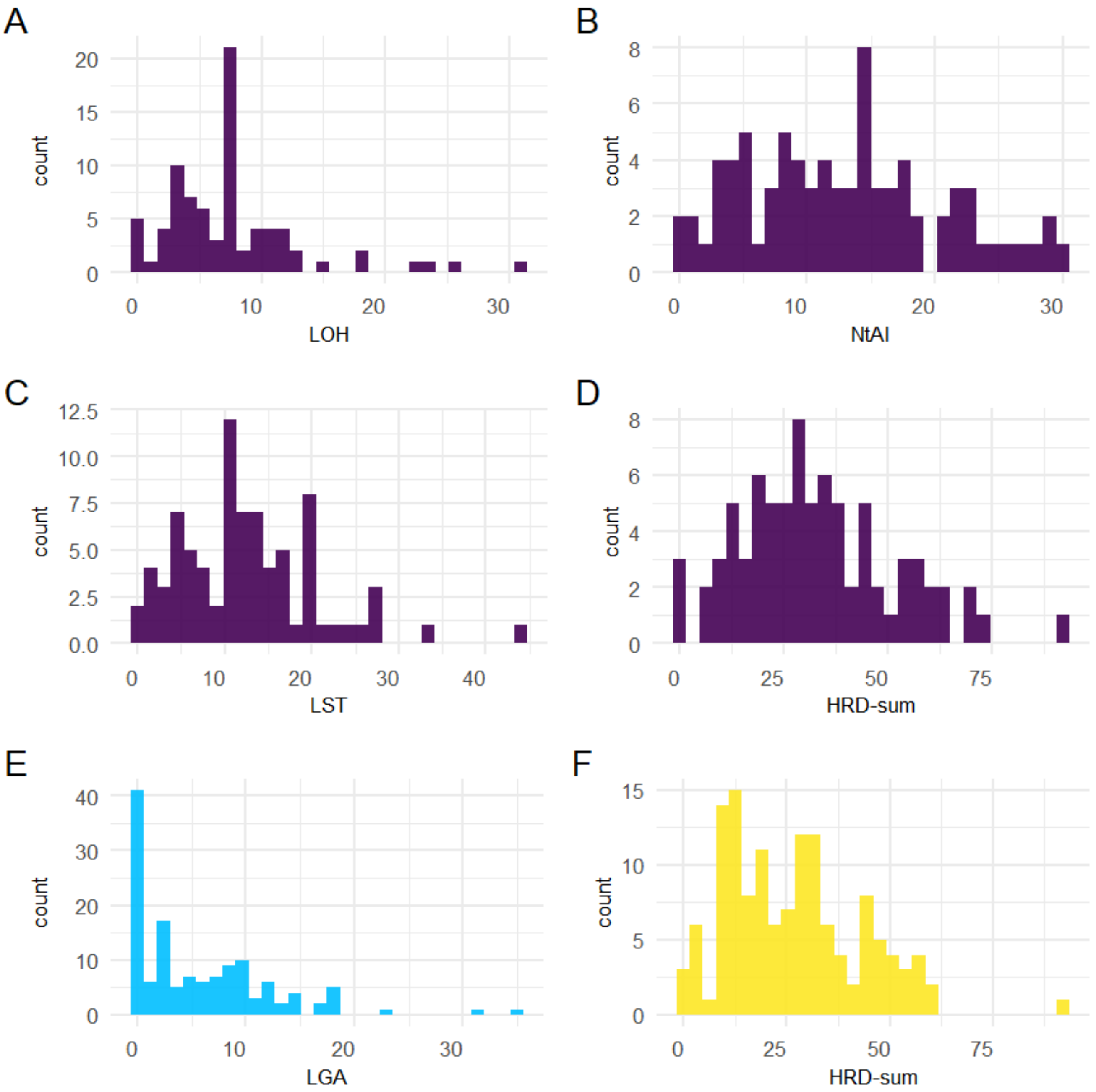

**Supplemental Figure 7.**

**Relationship between HRD-sum from WES and targeted panel (n=35). A,** HRD-sum determined by WES and targeted panel with the line of equality (blue). Pearson's coefficient= 0.69, p-value=5.4e-06. A symmetric distribution can be observed around the line of equality. **B,** Difference between measurements by WES and targeted panel for each subject against their mean. Gray line marks the zero and solid orange line indicates the mean difference, which is expected to be closer to zero. Dashed lines determine the 95% limits of agreement. Through this plot it is possible to explore any relationship between discrepancies (difference between measurements, y-axis) and the true value (determined in this setting as the mean between both methods). As expected, no correlation is found between difference and the mean (Spearman's rank correlation= 0.044). A random distribution around the difference can be seen throughout mean values. HRD-sum from WES have slightly higher values than panel, which can be seen in the mean difference >0 (orange solid line).

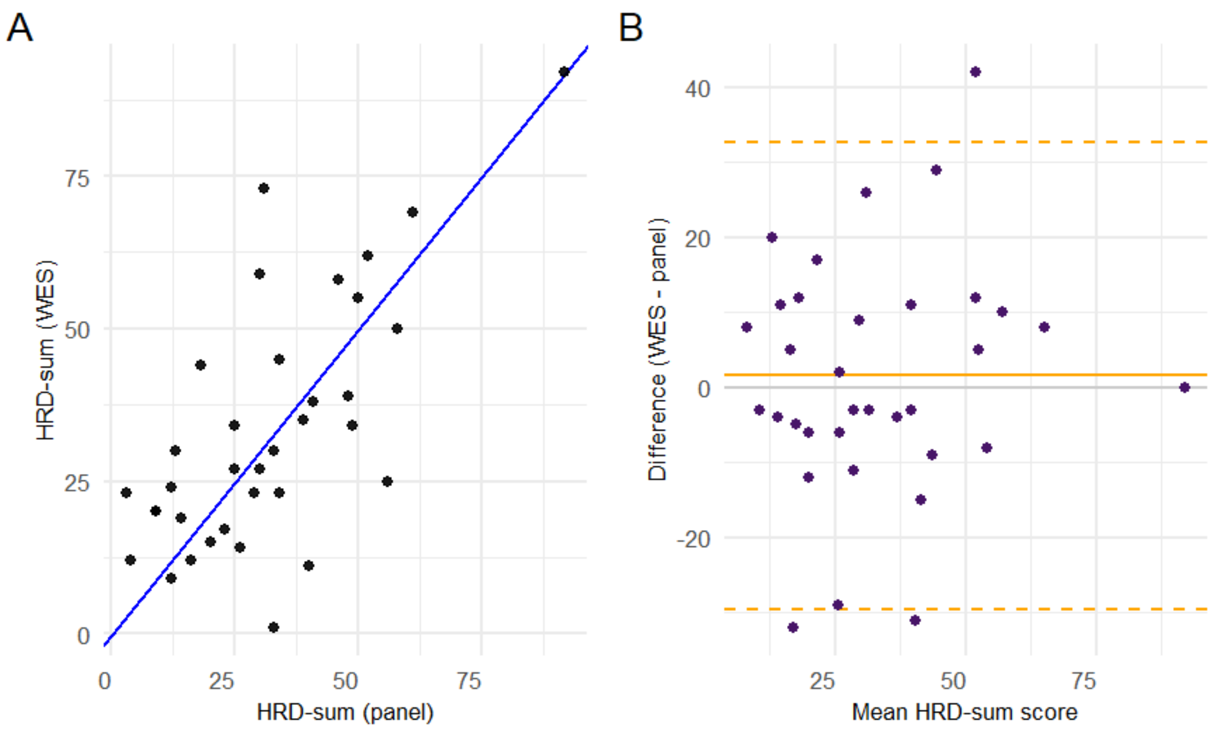

### Supplemental Figure 8.

**Relationship between scars and RAD51. A.** LGA from LP-WGS, **B.** LST from WES, **C.** LOH from WES, **D.** NtAI from WES.

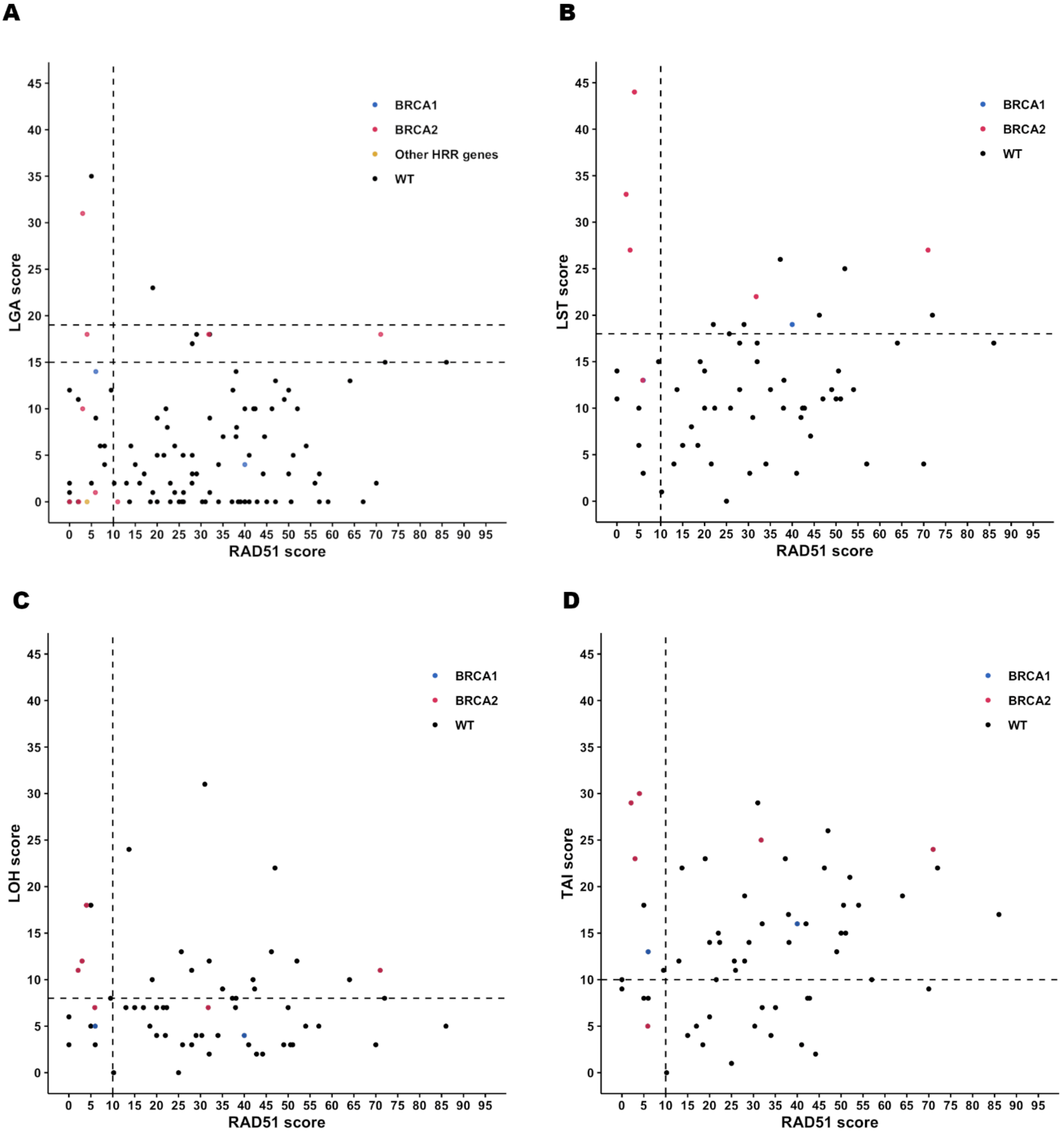

Supplemental Figure 9.

**Comparison between RAD51 low and high and each scar. A**. LOH, **B**. NtAI, **C.** LST, and **D.** the HRD-sum from WES, **E.** The LGA genomic scar from LP-WGS, **F.** The HRD-sum from targeted panel.

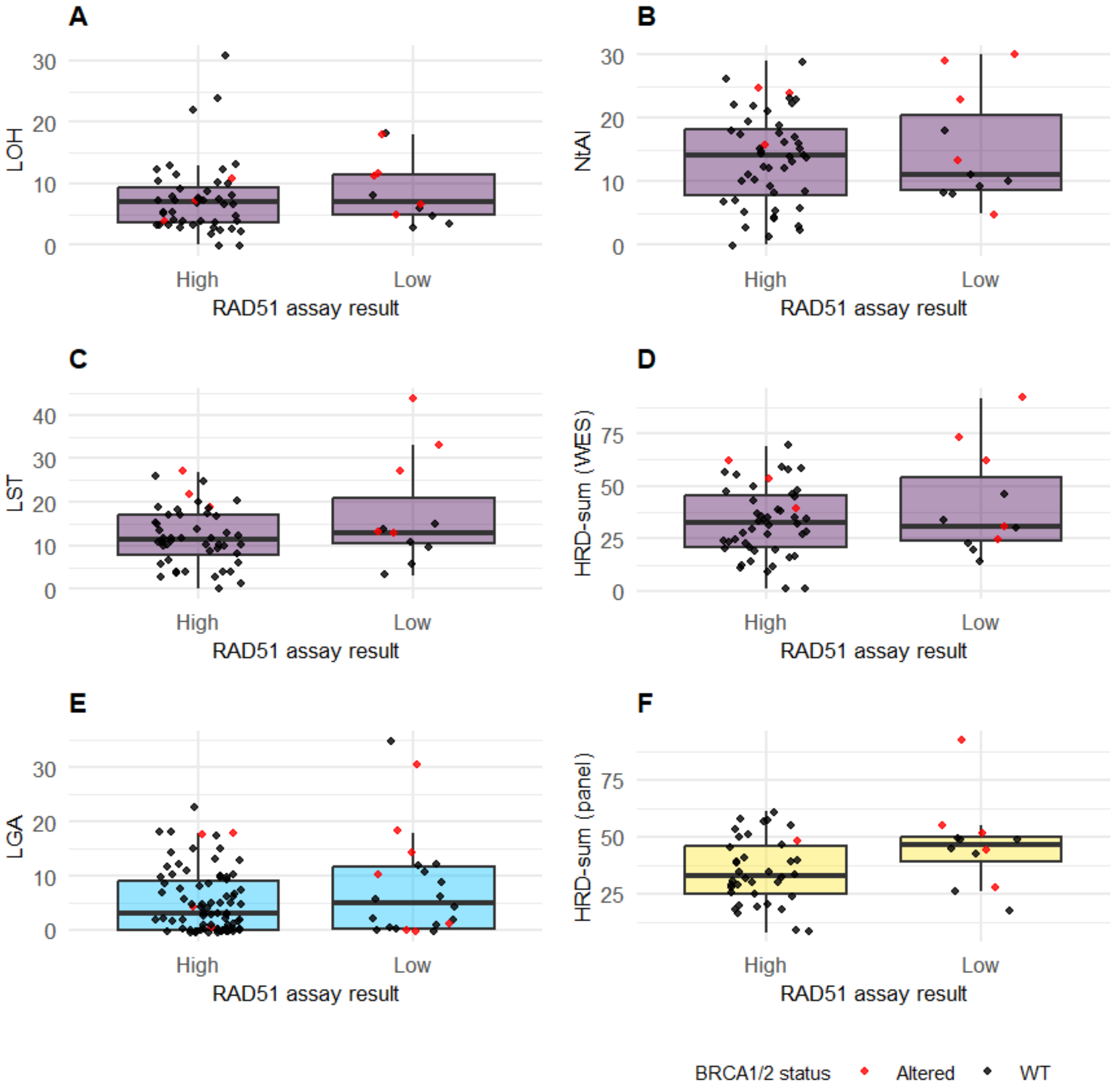

### Supplemental table 1.

| **Supp. Table 1. Curated list with genes of interest.** | | | | | |
|---|---|---|---|---|---|
| AR | KDM5A | ERCC5 | PALB2 | BRAF | PIK3CB |
| FOXA1 | KDM6A | BAP1 | RAD50 | EP300 | PTEN |
| SPOP | KMT2C | BARD1 | RAD51 | ERG | AKT1 |
| CCND1 | KMT2D | BLM | RAD51B | B2M | |
| CDK4 | PARP1 | BRCA1 | RAD51C | MYC | |
| CDK6 | MUTYH | BRCA2 | RAD51D | ARID1A | |
| CDKN2B | MLH1 | BRIP1 | ATM | MET | |
| MDM2 | MSH2 | CHEK1 | ATR | CTNNB1 | |
| RB1 | MSH6 | CHEK2 | PRKDC | APC | |
| TP53 | ERCC2 | FANCA | CDK12 | PIK3CA | |

### Supplemental table 2.

**Supp Table 2. Description of the BRCA1/2 altered cases.**

| Sample ID | Biopsy site | Prim / Met | Castration Sensitivity Status | Genomic Test | RAD51 score | BRCA1 | Alteration Type | BRCA1 LOH | Alteration state BRCA1 | BRCA2 | Alteration Type | BRCA2 LOH | Alteration state BRCA2 |
|---|---|---|---|---|---|---|---|---|---|---|---|---|---|
| **PRO205** | Prostate | Prim | HSPC | Targ. Seq | 0,00 | | | Yes* | WT | Frameshift Variant | NA | Yes | BIALLELIC |
| **PRO238** | Prostate | Prim | HSPC | Targ. Seq | 0,00 | | | No | WT | Frameshift Variant | Germinal* | Yes | BIALLELIC |
| **PRO277** | Prostate | Prim | HSPC | Targ. Seq | 0,00 | | | Yes | WT | HOMDEL | NA | NA | BIALLELIC |
| **PRO132** | Prostate | Prim | HSPC | Targ. Seq | 1,00 | | | No | WT | Frameshift Variant | Germinal* | Yes* | BIALLELIC* |
| **PRO043.2** | Bone | Met | CRPC | WES | 2,10 | | | No | WT | HOMDEL | Somatic | NA | BIALLELIC |
| **PRO113.1** | Lymph node | Met | CRPC | WES | 3,00 | | | No | WT | Frameshift Variant; Frameshift Variant | Germinal; Somatic | No | BIALLELIC |
| **PRO180.1** | Prostate | Prim | HSPC | Targ. Seq | 3,00 | | | No | WT | HOMDEL | NA | NA | BIALLELIC |
| **PRO153** | Liver | Met | CRPC | WES | 4,00 | | | No | WT | Frameshift Variant | Germinal | Yes | BIALLELIC |
| **PRO120** | Prostate | Prim | HSPC | WES | 5,90 | Frameshift Variant | Somatic | No | MONOALLELIC | Frameshift Variant | Germinal | Yes | BIALLELIC |
| **PRO055.2** | Prostate | Prim | HSPC | WES | 6,00 | Frameshift Variant | Somatic | No | MONOALLELIC | | | Yes | WT |
| **PRO084** | Prostate | Prim | HSPC | Targ. Seq | 11,00 | | | ND | WT | Nonsense Mutation | NA | ND | MONOALLELIC* |
| **PRO116** | Lymph node | Met | CRPC | WES | 31,77 | | | No | WT | Frameshift Variant | Somatic | Yes | BIALLELIC |
| **PRO055.1** | Liver | Met | HSPC | WES | 40,00 | Frameshift Variant | Somatic | No | MONOALLELIC | | | No | WT |
| **PRO014** | Liver | Met | CRPC | WES | 71,00 | | | No | WT | Nonsense Mutation | Germinal | Yes | BIALLELIC |

Abbreviations: Prim, Primary tumor; Met, Metastasis; HSPC, hormone-sensitive prostate cancer; CRPC, castration-resistant prostate cancer; Targ. Seq, Targeted panel sequencing; ND, not determined; NA, not applicable; HOMDEL, homozygous deletion; WT, wild type.

### Supplemental table 3.

**Supp Table 3. Cases with more than one sample from the same patient. Descriptive characteristics.**

| Patient ID | Collection date | Anatomic site | Prim/Met | Castration sensitivity status | RAD51 score | RAD51 result* | HRR gene alterations |
|---|---|---|---|---|---|---|---|
| **PRO018** | 31/07/2014 | Prostate | Prim | HSPC | 25 | High | WT |
| **PRO018** | 20/12/2017 | Lymph node | Met | CRPC | 6 | Low | WT |
| **PRO018** | 31/07/2020 | Lymph node | Met | CRPC | 50,56 | High | WT |
| **PRO028** | 08/09/2020 | Lymph node | Met | CRPC | 22,31 | High | WT |
| **PRO028** | 25/06/2004 | Prostate | Prim | HSPC | 30,3 | High | WT |
| **PRO054** | 19/04/2018 | Prostate | Prim | HSPC | 10,2 | High | WT |
| **PRO054** | 08/11/2019 | Lymph node | Met | CRPC | 25,92 | High | WT |
| **PRO055** | 09/01/2019 | Prostate | Prim | HSPC | 6 | Low | Altered |
| **PRO055** | 15/03/2019 | Liver | Met | HSPC | 40 | High | Altered |
| **PRO057** | 21/02/2019 | Prostate | Prim | HSPC | 49 | High | WT |
| **PRO057** | 22/03/2019 | Lymph node | Met | HSPC | 51,04 | High | WT |
| **PRO112** | 14/09/2020 | Bone | Met | HSPC | 21,53 | High | WT |
| **PRO112** | 17/06/2020 | Prostate | Prim | HSPC | 70 | High | WT |
| **PRO149** | 11/08/2021 | Bone | Met | CRPC | 86 | High | WT |
| **PRO149** | 21/08/2015 | Prostate | Prim | HSPC | 13 | High | WT |
| **PRO166** | 03/06/2021 | Prostate | Prim | HSPC | 15 | High | WT |
| **PRO166** | 16/03/2022 | Bone | Met | CRPC | 17 | High | WT |
| **PRO179** | 04/08/2021 | Bone | Met | HSPC | 52 | High | WT |
| **PRO179** | 10/08/2021 | Urethra | Prim | HSPC | 32 | High | WT |
| **PRO206** | 19/01/2022 | Bone | Met | CRPC | 29 | High | WT |
| **PRO206** | 04/04/2019 | Prostate | Prim | HSPC | 28 | High | WT |

REFERENCES: prim=primary, Met= metastasis, CRPC= castration-resistant prostate cancer, HSPC= hormone-sensitive prostate cancer, WT= wild-type.

* RAD51 result based on the 10% threshold

### Supplemental table 4.

**Supp Table 4. Regression analysis results for BRCA1/2 as response variable. Univariate and multivariate models for HRD-sum derived from targeted panel and WES.**

| Variable | Level | n | OR (95% IC) | p-value |
|---|---|---|---|---|
| **Targeted panel** | | | | |
| **HRD-sum (univariate)** | | 134 | 1.052 (1.018; 1.093) | 0.004 |
| **HRD-sum** | | 134 | 1.052 (1.018; 1.093) | 0.004 |
| **+ TP53** | Wild type | 79 | Ref | NA |
| | Altered | 55 | 0.608 (0.152; 2.081) | 0.445 |
| **HRD-sum** | | 134 | 1.061 (1.022; 1.106) | 0.003 |
| **+ Castration Sens. Status** | CRPC | 16 | Ref | NA |
| | HSPC | 118 | 2.413 (0.427; 23.467) | 0.374 |
| **HRD-sum** | | 134 | 1.058 (1.018; 1.105) | 0.006 |
| **+ Biopsy site** | Bone | 10 | Ref | NA |
| | Liver | 5 | 3.563 (0.178; 110.562) | 0.403 |
| | Lymph node | 12 | 0.867 (0.03; 25.294) | 0.925 |
| | Other | 6 | 0 (NA; 7.0115589128125e+43) | 0.992 |
| | Prostate | 101 | 1.931 (0.27; 40.28) | 0.573 |
| **HRD-sum** | | 134 | 1.058 (1.02; 1.102) | 0.004 |
| **+ Prim.Met** | Met | 30 | Ref | NA |
| | Prim | 104 | 1.589 (0.397; 7.802) | 0.534 |
| **HRD-sum** | | 122 | 1.054 (1.018; 1.097) | 0.005 |
| **+ Grade Group categorized** | 1-3 | 25 | Ref | NA |
| | 4-5 | 97 | 1.986 (0.324; 38.301) | 0.533 |
| **HRD-sum** | | 133 | 1.053 (1.018; 1.094) | 0.004 |
| **+ Stage Metastasis** | M0 | 49 | Ref | NA |
| | M1 | 84 | 0.823 (0.239; 3.081) | 0.761 |
| **WES** | | | | |
| **HRD-sum (univariate)** | | 80 | 1.072 (1.031; 1.127) | 0.002 |
| **HRD-sum** | | 80 | 1.077 (1.033; 1.137) | 0.002 |
| **+ TP53** | Wild type | 48 | Ref | NA |
| | Altered | 32 | 2.05 (0.445; 10.798) | 0.364 |
| **HRD-sum** | | 80 | 1.068 (1.025; 1.125) | 0.004 |
| **+ Castration Sens. Status** | CRPC | 35 | Ref | NA |
| | HSPC | 45 | 0.624 (0.112; 3.077) | 0.564 |
| **HRD-sum** | | 80 | 1.076 (1.029; 1.138) | 0.004 |
| **+ Biopsy site** | Bone | 24 | Ref | NA |
| | Liver | 8 | 4.826 (0.449; 60.887) | 0.192 |
| | Lymph node | 15 | 3.889 (0.473; 42.223) | 0.217 |
| | Other | 3 | 0 (NA; 3.81356723836726e+70) | 0.994 |
| | Prostate | 30 | 1.726 (0.161; 20.616) | 0.642 |
| **HRD-sum** | | 80 | 1.07 (1.027; 1.126) | 0.003 |
| **+ Prim.Met** | Met | 48 | Ref | NA |
| | Prim | 32 | 0.571 (0.075; 3.018) | 0.533 |
| **HRD-sum** | | 70 | 1.066 (1.024; 1.12) | 0.004 |
| **+ Grade Group categorized** | 1-3 | 9 | Ref | NA |
| | 4-5 | 61 | 8181985.683 (0; NA) | 0.994 |
| **HRD-sum** | | 79 | 1.078 (1.034; 1.133) | 0.001 |
| **+ Stage Metastasis** | M0 | 24 | Ref | NA |
| | M1 | 55 | 2.324 (0.458; 16.523) | 0.344 |

### Supplemental table 5.

**Supp Table 5. Regression analysis results for RAD51 as response variable. Univariate and multivariate models for HRD-sum derived from targeted panel and WES.**

| Variable | Level | n | OR (95% IC) | p-value |
|---|---|---|---|---|
| **Targeted panel** | | | | |
| **HRD-sum** | | 99 | 1.033 (1.006; 1.064) | 0.021 |
| **HRD-sum** | | 99 | 1.034 (1.006; 1.064) | 0.02 |
| **+ TP53** | Wild type | 56 | Ref | NA |
| | Altered | 43 | 0.826 (0.308; 2.145) | 0.697 |
| **HRD-sum** | | 99 | 1.03 (1.001; 1.062) | 0.049 |
| **+ Castration Sens. Status** | CRPC | 12 | Ref | NA |
| | HSPC | 87 | 0.647 (0.167; 2.745) | 0.534 |
| **HRD-sum** | | 99 | 1.035 (1.002; 1.069) | 0.037 |
| **+ Biopsy site** | Bone | 4 | Ref | NA |
| | Liver | 5 | 0.053 (0.001; 1.05) | 0.082 |
| | Lymph node | 11 | 0.22 (0.009; 2.476) | 0.254 |
| | Other | 5 | 0.072 (0.002; 1.298) | 0.106 |
| | Prostate | 74 | 0.147 (0.007; 1.327) | 0.116 |
| **HRD-sum** | | 99 | 1.026 (0.996; 1.059) | 0.097 |
| **+ Prim.Met** | Met | 22 | Ref | NA |
| | Prim | 77 | 0.542 (0.174; 1.739) | 0.293 |
| **HRD-sum** | | 91 | 1.035 (1.006; 1.068) | 0.021 |
| **+ Grade Group categorized** | 1-3 | 15 | Ref | NA |
| | 4-5 | 76 | 0.565 (0.154; 2.349) | 0.4 |
| **HRD-sum** | | 98 | 1.033 (1.005; 1.066) | 0.027 |
| **+ Stage Metastasis** | M0 | 29 | Ref | NA |
| | M1 | 69 | 0.372 (0.135; 1.016) | 0.053 |
| **WES** | | | | |
| **HRD-sum** | | 59 | 1.023 (0.988; 1.061) | 0.204 |
| **HRD-sum** | | 59 | 1.023 (0.987; 1.062) | 0.205 |
| **+ TP53** | Wild type | 34 | Ref | NA |
| | Altered | 25 | 1.162 (0.293; 4.497) | 0.826 |
| **HRD-sum** | | 59 | 1.034 (0.994; 1.078) | 0.104 |
| **+ Castration Sens. Status** | CRPC | 25 | Ref | NA |
| | HSPC | 34 | 2.442 (0.535; 13.614) | 0.269 |
| **HRD-sum** | | 59 | 1.052 (1.006; 1.108) | 0.035 |
| **+ Biopsy site** | Bone | 11 | Ref | NA |
| | Liver | 8 | 0.389 (0.012; 6.098) | 0.523 |
| | Lymph node | 13 | 1.098 (0.104; 12.038) | 0.935 |
| | Other | 3 | 0 (NA; 5.85660524257388e+67) | 0.994 |
| | Prostate | 24 | 4.138 (0.57; 48.268) | 0.197 |
| **HRD-sum** | | 59 | 1.035 (0.996; 1.08) | 0.084 |
| **Prim.Met** | Met | 33 | Ref | NA |
| | Prim | 26 | 3.006 (0.672; 15.689) | 0.162 |
| **HRD-sum** | | 54 | 1.026 (0.99; 1.066) | 0.158 |
| **+ Grade Group categorized** | 1-3 | 4 | Ref | NA |
| | 4-5 | 50 | 0.552 (0.058; 12.177) | 0.631 |
| **HRD-sum** | | 59 | 1.014 (0.977; 1.054) | 0.45 |
| **+ Stage Metastasis** | M0 | 19 | Ref | NA |
| | M1 | 40 | 0.368 (0.085; 1.554) | 0.169 |
